# PANDORA v2.0: Benchmarking peptide-MHC II models and software improvements

**DOI:** 10.1101/2023.07.20.549892

**Authors:** Farzaneh M. Parizi, Dario F. Marzella, Gayatri Ramakrishnan, Peter A. C. ‘t Hoen, Mohammad Hossein Karimi-Jafari, Li C Xue

## Abstract

T-cell specificity to differentiate between self and non-self relies on T-cell receptor (TCR) recognition of peptides presented by the Major Histocompatibility Complex (MHC). Investigations into the three-dimensional (3D) structures of peptide:MHC (pMHC) complexes have provided valuable insights of MHC functions. Given the limited availability of experimental pMHC structures and considerable diversity of peptides and MHC alleles, it calls for the development of efficient and reliable computational approaches for modeling pMHC structures. Here we present an update of PANDORA and the systematic evaluation of its performance in modelling 3D structures of pMHC class II complexes (pMHC-II), which play a key role in the cancer immune response. PANDORA is a modelling software that can build low-energy models in a few minutes by restraining peptide residues inside the MHC-II binding groove. We benchmarked PANDORA on 136 experimentally determined pMHC-II structures covering 44 unique αβ chain pairs. Our pipeline achieves a median backbone Ligand-Root Mean Squared Deviation (L-RMSD) of 0.42 Å on the binding core and 0.88 Å on the whole peptide for the benchmark dataset. We incorporated software improvements to make PANDORA a pan-allele framework and improved the user interface and software quality. Its computational efficiency allows enriching the wealth of pMHC binding affinity and mass spectrometry data with 3D models. These models can be used as a starting point for molecular dynamics simulations or structure-boosted deep learning algorithms to identify MHC-binding peptides. PANDORA is available as a Python package through Conda or as a source installation at https://github.com/X-lab-3D/PANDORA.

## Introduction

The ability of T-cells to recognize and eliminate infected or transformed cells relies on their ability to distinguish between self and non-self peptides presented by the Major Histocompatibility Complex (MHC) on the surface of these cells. Upon recognition of a non-self peptide by T-cell receptors (TCR), T-cells activate and initiate an immune response. MHC class I (MHC-I) molecules typically present intracellular antigens to cytotoxic CD8+ T-cells, which eliminate the cell presenting the antigen. MHC-II molecules present extracellular antigens to helper CD4+ T-cells, which assist other immune cells by releasing cytokines and orchestrating the immune response (1,2). To unravel the mechanisms of peptide presentation to T-cells and immune response, it is essential to investigate how peptides bind to MHC molecules.

Understanding the mechanism of peptide-MHC (pMHC) binding raises an intriguing research question regarding how MHC molecules effectively bind to a wide range of peptides while maintaining strong binding and specificity. Previous research focusing on the structural aspects of pMHC complexes has provided valuable insights into our understanding of antigen presentation specificity (3) and peptide binding dynamics (4,5). Allele-specific residues at anchor positions and complementary pockets in the MHC molecule play a significant role in determining the promiscuity and specificity of peptide recognition by MHC molecules (6,7). Notably, the presence of hydrophobic anchors and the formation of hydrogen bonds have been discovered to stabilize the pMHC-II interaction (8,9). Similarly, in the case of MHC class I, peptide-dependent stability is achieved through the establishment of conserved hydrogen bonds at the N and C termini of peptides, along with anchor residues that fit into pockets of MHC class I (10,11). Furthermore, structural investigations have provided insights into other mechanisms, such as the molecular basis of autoimmune diseases (10) and T-cell recognition (11,12). The knowledge gained from structural studies has also facilitated the design of novel therapies and can help the development of effective vaccine strategies (13,14). Therefore, access to structural information on pMHC is crucial for these advancements.

This work focuses on pMHC-II binding. MHC-II is crucial in antigen presentation, particularly for extracellular antigens. Additionally, MHC-II mediated CD4+ T-cell responses are reported to account for the predominant immune responses following cancer vaccine treatment (15–18). The MHC-II complex consists of two membrane-anchored chains: an α- and β-chain (Fig. 1), and it can bind peptides up to 25 residues in length (19,20). The binding groove of MHC-II can hold a 9-mer core (21). The residues outside the groove form the Peptide Flanking Regions (PFR), namely the left (N-terminus) and right (C-terminus) PFRs. A peptide is kept in place within the groove by three or four main conserved binding pockets: Pockets 1, 4, 6, and 9 (Fig. 1B), and alongside these, there are smaller auxiliary anchor pockets (22,23).

**Figure 1:**
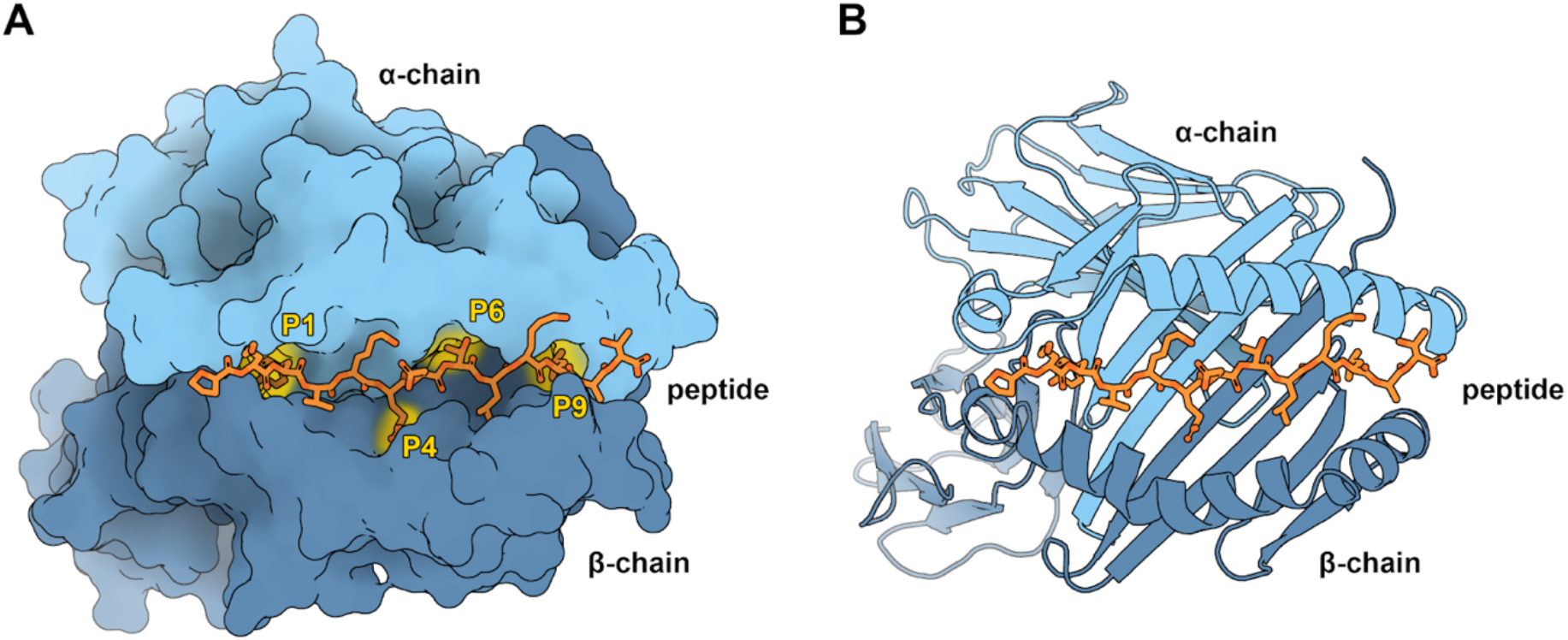
Overview of the pMHC-II complex. (A) Representation of an MHC-II molecule by its accessible surface area, visualized with Protein Imager (24). MHC-II consists of an α-chain (light blue) and a β-chain (dark blue). Shown are the four characteristic pockets in the binding groove (P1, P4, P6, and P9), occupied by the corresponding peptide (orange) anchor residues. As modeling restraints (yellow), PANDORA uses the atomic contacts between the peptide anchor residues and the MHC-II pockets. (B) Cartoon representation of pMHC-II. The peptide binding groove consists of two α-helices on a floor of β-sheets, in which the peptide resides (PDB ID: 1DLH).

To accommodate a diverse range of antigens within the MHC groove, the MHC locus stands out as the most polymorphic region in the human genome (2,25). With over 10,754 alleles for MHC class II, there is a significant variation in MHC-II alleles and the peptides they can bind (26). Unfortunately, only a few pMHC-II structures have been experimentally resolved (about 240 entries in the PDB, the Protein Data Bank (27)). This necessitates the development of fast, structure-based computational modeling methods to overcome the scarcity of available pMHC-II structures. However, only a few modeling methods have been explicitly developed for pMHC-II complexes.

Most existing pMHC-II modelling methods rely on grid-based docking, including pDock and EpiDock (28–33). Among them, pDock has demonstrated improved performance in generating peptide core conformations bound to MHC-II. The pDock’s approach involves receptor modeling followed by flexible peptide docking into the binding groove while retaining its starting conformation using loose restraints. Current pMHC-II modeling approaches are often limited in terms of usability due to: 1) long computation times; 2) the use of closed-source software; 3) limited coverage of diverse MHC alleles; and 4) uncertainty regarding the quality of PFR conformations. Additionally, structural modelling of pMHC-II complexes is fraught with challenges. It is not always clear which region of a peptide forms the core and is directly anchored to the MHC-II receptors (34,35). Existing methods, such as NetMHCIIpan-4.0 (36), can provide reasonably accurate predictions for the binding core. Furthermore, the flexibility of PFRs poses additional hurdles. To address these challenges, the development of fast and pan-allelic pMHC-II modelling software is required to integrate prediction of the binding core and generation of plausible conformations for the entire peptide bound to MHC-II.

We present here the utility and performance of our pMHC modelling software, PANDORA v2.0, for pMHC-II modeling and its new version updates. We have earlier demonstrated PANDORA’s reliable performance for modeling pMHC-I complexes (37,38). PANDORA leverages two pieces of domain knowledge: 1) the high conservation of MHC structures and 2) the anchoring of peptides to the main pockets of MHC molecules (**Fig. 2**). We benchmarked PANDORA on 136 experimentally resolved pMHC-II structures. When compared with an existing pMHC-II modelling technique, pDock (32), and also with AlphaFold (39), we show that PANDORA outperforms these methods in terms of generated model quality and computational efficiency. Additionally, we evaluate the effectiveness of the anchor prediction tool used in our approach (NetMHCIIpan-4.0). PANDORA’s quality and speed show the potential for boosting structure-based Deep Learning (DL) algorithms, making it a valuable tool in developing effective vaccine designs. We also discuss the existing limitations of anchor predictions and propose the integration of a structural and physics-based anchor predictor as a potential solution. Furthermore, we highlight the importance of further research in the modeling of post-translational modifications (PTMs) on peptide-MHC interactions.

**Figure 2:**
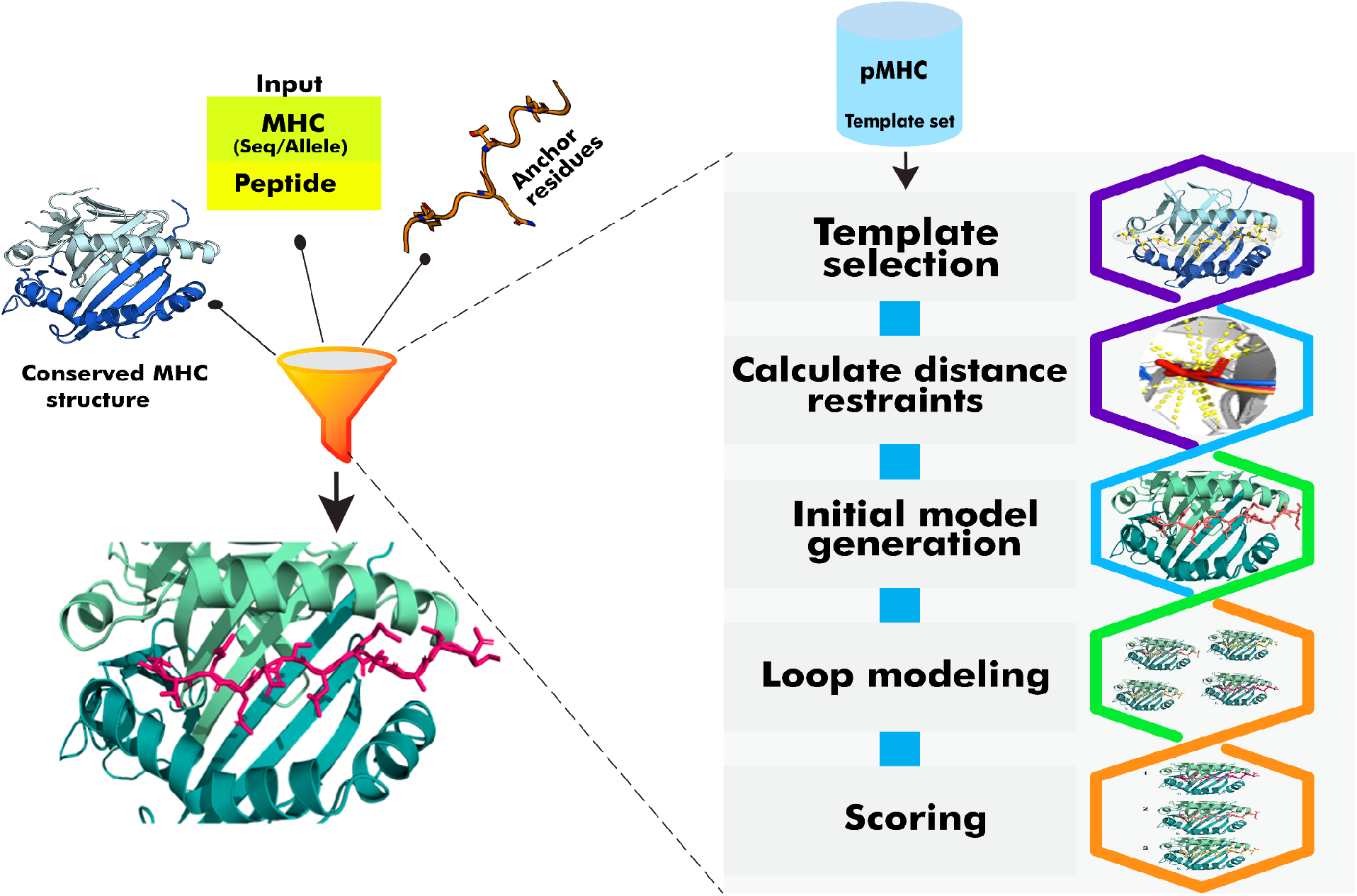
Overview of the PANDORA pMHC modelling framework. PANDORA as an integrative modelling protocol, leverages two domain knowledge aspects: the highly conserved nature of MHC structures and the binding of peptides to MHC pockets with anchor residues. PANDORA takes the sequence information of a target peptide and MHC as input and selects a template pMHC structure from a template set based on sequence similarity. The target peptide core is superposed onto the template peptide core. In flexible mode, it applies distance restraints for anchor residues. The framework performs loop modeling of flexible regions and energy minimization of pMHC-II conformations. Conformations are ranked to select those resembling the near-native conformation.

## Materials and Methods

### Structural template set building

Building the template dataset is similar to our previous work (37) and is expanded to make it suitable for pMHC-II. PANDORA retrieved structures of pMHC-II complexes from IMGT/3Dstructure-DB (40) and filters for those with peptides of lengths between 7 and 25 residues. Structures including the DM chaperone and the CLIP peptide, both known to affect the MHC-II conformation, are discarded (20,41). The MHC-II alpha chain is renamed as chain “M” and the beta chain is renamed as chain “N” to make a distinction from MHC-I β2-microglobulin which is renamed as chain “B”. The peptide chain is renamed as chain “P”. For the benchmark experiment presented in this work, the parsing resulted in a total of 136 pMHC-II templates, spanning over 32 α chain alleles, 81 β chain alleles, and a total of 44 unique MHC-II αβ pairs (see details in Supplementary Table S1).

### BLAST databases generation

BLAST (v2.10) is used to assign allele names (needed by NetMHCIIpan-4.0 for predicting binding cores) to the MHC sequences provided by the user and, independently, for the template selection step. The current version of PANDORA uses two BLAST databases. The first one (BLAST-DB1) is generated from the manually curated MHC sequences taken from https://www.ebi.ac.uk/ipd/, and it is used to assign the allele name to any MHC sequence provided by the user. This allele name will later be used as input for NetMHCIIpan4.0 to predict the binding core (see Template selection). The second one (BLAST-DB2) is generated from the template set sequences extracted by the PDB files retrieved as described above, and it is used for the template selection step.

### Template selection

The template selection step has been updated from the first version of PANDORA (which used allele type names to identify templates) to a BLAST-based template selection. First, the target MHC sequences are queried against the BLAST-DB2 database with default parameters, and the results are ranked by percentage sequence identities. Templates sharing the highest sequence identity with the target sequences are selected and further ranked by peptide alignment score. Our peptide alignment method includes alignment of the binding core of the peptides followed by the addition of gaps at both their termini to account for different peptide lengths. The binding cores of the templates are derived from their corresponding structures. The binding core for the query peptide is predicted by NetMHCIIpan4.0. The peptides’ alignments are then scored using a PAM30 substitution matrix. The highest-ranking template is then selected for modeling.

### Modeling

We perform 3D modeling as described previously. For MHC-II, we restrain four anchor positions (P1, 4, 6, and 9) while keeping the peptide flanking regions flexible during the modeling step. In the default mode for pMHC-II cases, the whole peptide core is kept fixed as the template conformation. PANDORA v2.0 also supports restraints-flexible modelling mode for the peptide core, where users can provide anchors’ restraints standard deviation, thereby specifying the extent of deviation of restraints from those in the templates in Angstroms. By default, 20 (adjustable) 3D models are produced, which are ranked by MODELLER’s (42) internal molpdf score.

### L-RMSD calculation

The L-RMSD is calculated as described in (43) as the backbone L-RMSD (including only the backbone atoms N, Cα, C, and O). We calculate “Core L-RMSD” for the binding core residues of the peptide, “Flanking L-RMSD” for the flanking regions of the peptide (i.e., the residues at the N-terminal of the first anchor and at the C-terminal of the fourth anchor), and “Whole L-RMSD” for all the residues of the peptide. The lower the L-RMSD, the better a model is.

## Results

### Modeling Performance on the Benchmark Set

We benchmarked PANDORA’s performance in reproducing X-ray crystal structures of pMHC-II complexes from the template set (n = 136). We carried out a leave-one-out validation approach where we iteratively removed a structure from the template database and allowed PANDORA to predict the pMHC-II complex using sequence and anchor information. To rule out the impact of anchor predictions, the anchor positions provided to PANDORA in this experiment were obtained from the target experimental structure to assess the modelling quality (see discussion on anchor prediction effects in the “NetMHCIIpan’s anchor prediction” section).

We analyzed the distribution of the best L-RMSD (i.e., the lowest L-RMSD) conformations obtained for the whole and core peptide regions (Fig. 3A,C,E; detailed information on different RMSD values is reported in Supplementary Table S2). The results demonstrate that for 91.1% (125 out of 136) cases, PANDORA was able to sample at least one high-quality model (whole peptide L-RMSD < 2 Å) with an overall mean L-RMSD of 1.11 ± 0.86 Å. A small number of cases (11 out of 136) showed a relatively higher whole peptide L-RMSD of > 2 Å (see Fig. 3D, F, and “The PFR Conformation Evaluation” section). We investigated the distribution of whole and core L-RMSDs over various peptide lengths, as illustrated in Figure 3A. Our analysis reveals a correlation between peptide lengths and the L-RMSD values, with longer peptides exhibiting higher L-RMSD values (Supplementary Fig. S1A–B).

**Figure 3:**
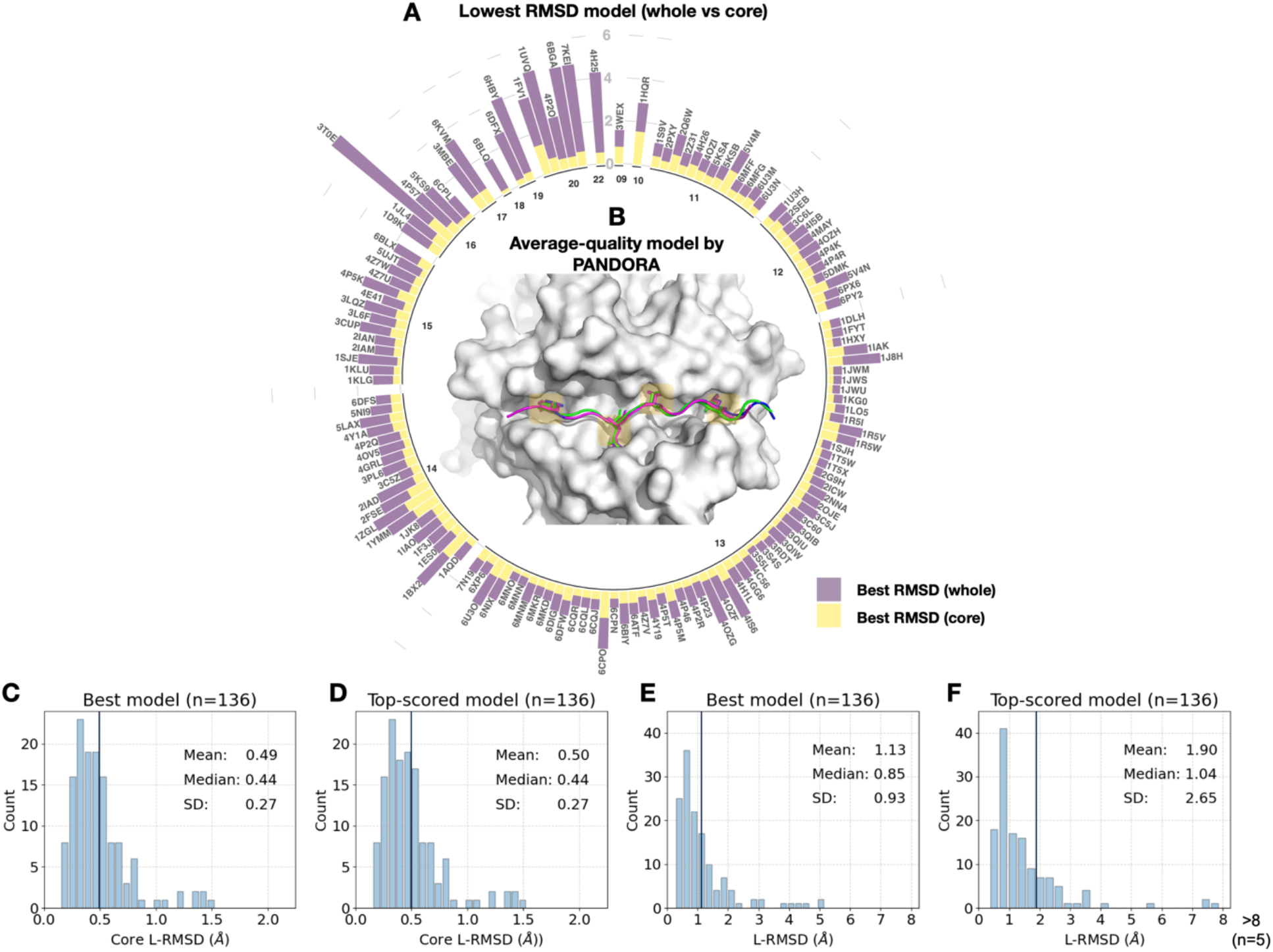
Benchmark results on reproducing 136 pMHC-II complexes with X-ray structures. (A) Sampling performance of the PANDORA benchmark experiment. A circular bar plot grouped based on peptide length (represented by the numbers in the inner circle) reports the lowest backbone L-RMSD (Y-axis) for the whole peptide (yellow) and binding core (purple). (B) An example of an average-quality 3D model generated by PANDORA. The target peptide (PDB ID: 4I5B) is marked in green; the template structure (PDB ID: 2OJE) is marked in magenta; and the PANDORA model structure (best conformation among the top 5 ranked) is marked in darkblue. (C, D) Histogram of the lowest backbone L-RMSD models in the peptide binding core vs. the whole peptide. (E, F) Complete performance of PANDORA (modeling + scoring). Histogram for the top-ranked models by PANDORA in terms of backbone L-RMSD on the peptide binding core vs. the whole peptide.

Furthermore, in terms of model ranking, we examined the performance of PANDORA by reporting L-RMSD for the best-scored model (conformation ranked as the top model using molpdf scoring) (Fig. 3E-F; for details, see Supplementary Fig. S2). Our results show that PANDORA achieved an 85% success rate (L-RMSD < 2 Å) for the top 5 ranked models in the entire template set.

### PANDORA generates low-energy conformations for the binding core

With four anchor positions in the binding groove, the structure of pMHC-II is well-suited for a restraint-based modelling approach. With the default mode (see Modelings in Methods), PANDORA demonstrates high accuracy in reproducing high-quality core conformations, with an average core L-RMSD of 0.49 ± 0.27 Å (93.38% of the cases having an L-RMSD < 1 Å) (Fig. 2C, 2E). The fully-flexible mode, which allows for flexibility in the peptides’ binding core, yielded an average core L-RMSD of 0.47 ± 0.2 Å (Supplementary Fig. S2). However, the restraints-flexible mode increases the computational time by 90%, while marginally enhancing the overall quality (∼ 6.46 min/case in the fully flexible mode vs. ∼ min/case in the default mode).

### Comparisons with AlphaFold and pDock

We compared PANDORA’s performance against existing approaches, such as pDOCK (32) and AlphaFold (39). To assess the general performance of the pipeline, we used NetMHCIIpan’s predicted anchor positions for this comparison.

pDock uses the ICM (Internal Coordinate Mechanics) algorithm to perform a flexible peptide docking into the MHC binding groove. During docking, the position of the peptide is only loosely constrained so that it retains a conformation close to its initial structure. For comparisons against pDock, we modeled pMHC-II complexes using PANDORA for the cases reported by Khan and Ranganathan (32). We obtained a mean L-RMSD of 0.27 ± 0.07 Å for Cα core while pDock achieved 0.59 ± 0.24 Å (Table 1). pDock retained RMSD estimates by redocking experimental pMHC X-ray structures; thus, the core residues are referred to as a priori. But, PANDORA automatically predicts anchor residues (using NetMHCIIpan-4.0(36)) and a suitable template, generating higher-quality peptide core conformations. We did not use pDock to perform cross-docking on our template set since pDock is not publicly available for download and usage.

**Table 1.**
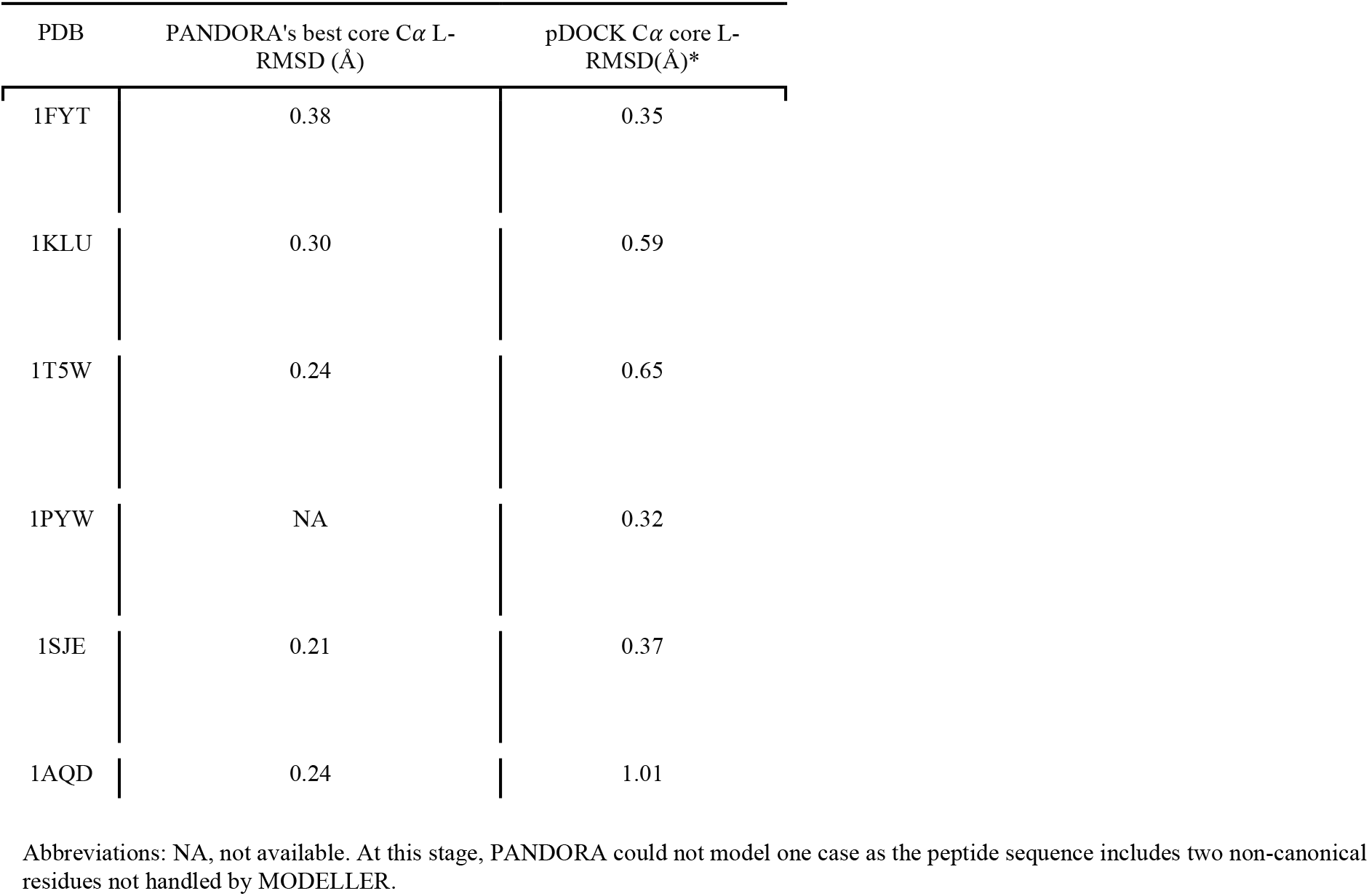
Comparison of PANDORA and pDock in pMHC-II modelling. The calculated core Cα L-RMSD (Å) on modeling 5 pMHC-II complexes using integrative homology modeling and a grid-based docking method. PANDORA’s best model quality is compared to pDock as no pDock scoring function was disclosed in pDock so it seems that pDock reported the best RMSDs in their paper. *: Data extracted from Khan & Ranganathan (32).

We also compared PANDORA with one of the best AI methods available for protein structure predictions, i.e., AlphaFold. AlphaFold is an advanced deep neural network approach that achieves unprecedented accuracy in protein folding predictions (44). However, since AlphaFold relies on sequence conservation information, it performs poorly on proteins where such information is absent, such as antibody-antigens and peptides (e.g., synthetic peptides or frame-shift mutated peptides) (45). For an objective comparison, we chose to use a version of AlphaFold that also uses templates to predict MHC structures (colabfold(44)). Our comparison shows that not all Alphafold-generated pMHC-II conformations have the correct anchor positions. Out of four randomly selected cases, in two cases (PDB ID: 3C5Z and 6PXP), AlphaFold was unsuccessful in predicting the peptide’s conformation with the correct anchor residues (Fig. 4A, C). Notably, they were both part of AlphaFold’s training set. This is mainly because PANDORA correctly identified the binding core for the four cases using NetMHCIIpan binding core predictions (see “NetMHCIIpan4.0 performance” in the next section).

**Figure 4:**
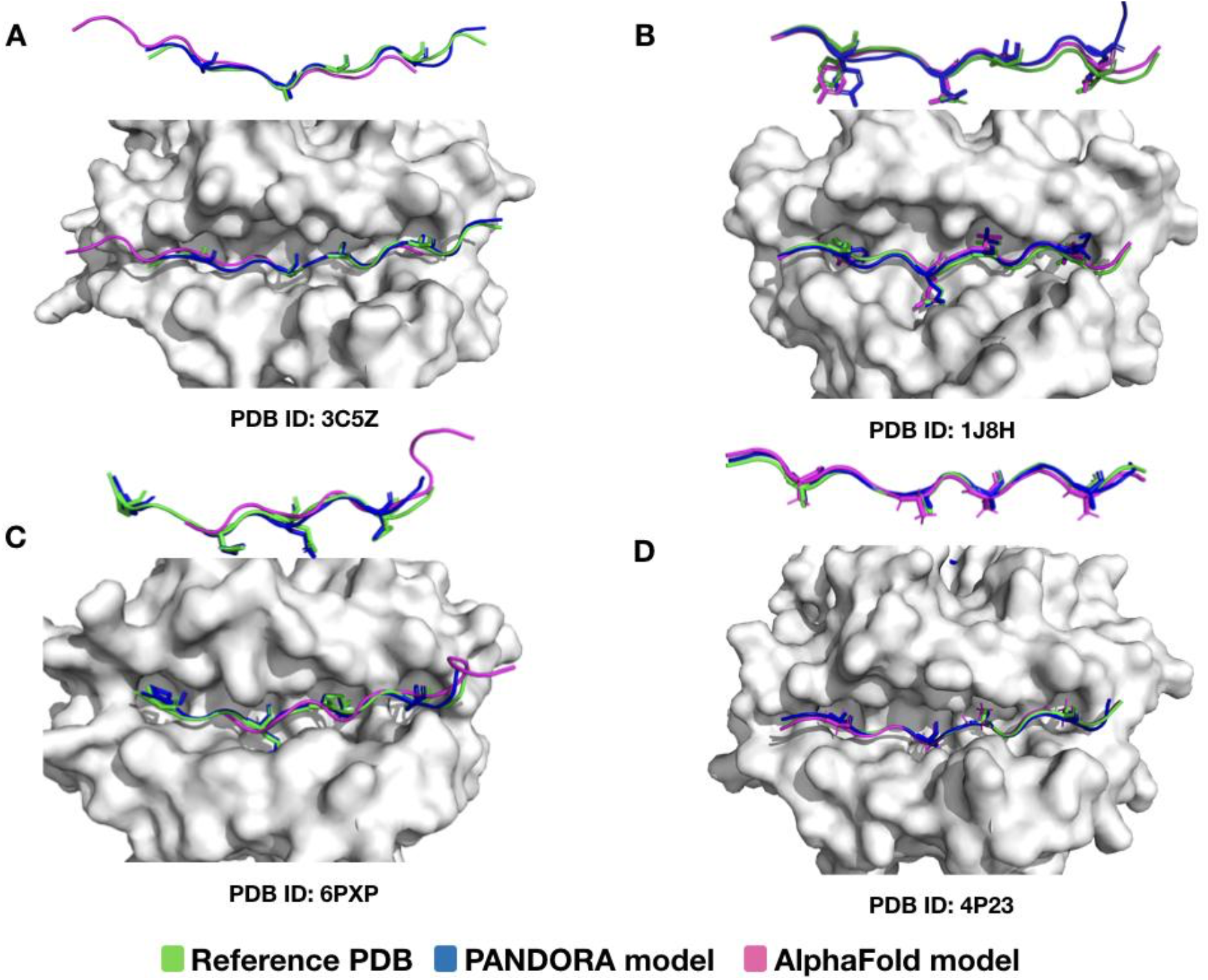
Comparison of PANDORA and AlphaFold in reproducing pMHC-II complexes. The peptide conformations are colored as follows: reference PDB in green, AlphaFold model in magenta, and PANDORA model in blue. The presented model for AlphaFold and PANDORA is the best RMSD model among the top-5 ranked models. Overall, AlphaFold models have quite good backbone predictions (probably due to the usage of templates), but in 2 out of 4 cases, the peptides are shifted. A) 24-mer peptide binding to H2-AB1*01 (PDB ID: 3C5Z); however, the predicted conformation is shifted by 3 residues; B) 13-mer peptide binding to HLA-DRA*0101, HLA-DRB1*0401 (PDB ID: 1J8H); C) 12-mer peptide binding to HLA-DQA1*0201, HLA-DQB1*0201 (PDB ID: 6PX6); however, the predicted AlphaFold binding core conformation is shifted by 4 residues; D) 13-mer peptide binding to H2-AB1*01 (PDB ID: 4P23) and the binding core is accurately identified by AlphaFold. Only A, B, and D are reportedly known to be present in AlphaFold training.

Additionally, considering computational cost, PANDORA outperforms AlphaFold (regarding resources) and pDock. PANDORA is much more efficient considering template selection, anchor prediction, and modeling require ∼3-4 minutes (from 3.75 to 6.46 minutes per case, depending on the mode, with shorter times for pMHC-I) on one core from an Intel(R) Xeon(R) Gold 6142 CPU @ 2.60 GHz. While pDock reported requiring 10 minutes for modeling on 2 CPUs 3.20 GHz (without homology modelling). Also, AlphaFold requires a significant amount of computation power-up to 18 GB of GPU power and 20 minutes to model a single pMHC case.

### The impacts of binding core prediction on PANDORA model quality

The interaction between the peptide binding core and the MHC binding groove directly impacts the quality of a model; therefore, choosing the correct binding core is critical. In the absence of user-defined anchor residues, PANDORA uses NetMHCIIpan to predict the binding core.

Hence, we evaluate NetMHCIIpan’s binding core prediction accuracy by comparing its predictions to known cores from experimental PDB structures. Our results show that the anchors were incorrectly predicted in 33 of the 136 cases in the benchmark dataset. In most cases, the observed shifts were by one or two residues (26 of 33), but misalignments of up to 8 residues were also observed (Supplementary Figure S4).

### PANDORA as a pan-allele modelling method

Owing to the high structural similarity across MHC-II alleles, it is possible to model pMHC-II complexes using different MHC-II alleles as templates. Our results show that even when a template with the same MHC allele type for either of the chains was not available in the template set (25% of cases), PANDORA was still able to provide models with a mean L-RMSD of 0.86 Å for the best-RMSD models and 1.05 Å for the top 10 ranked models (Fig. 5).

**Figure 5:**
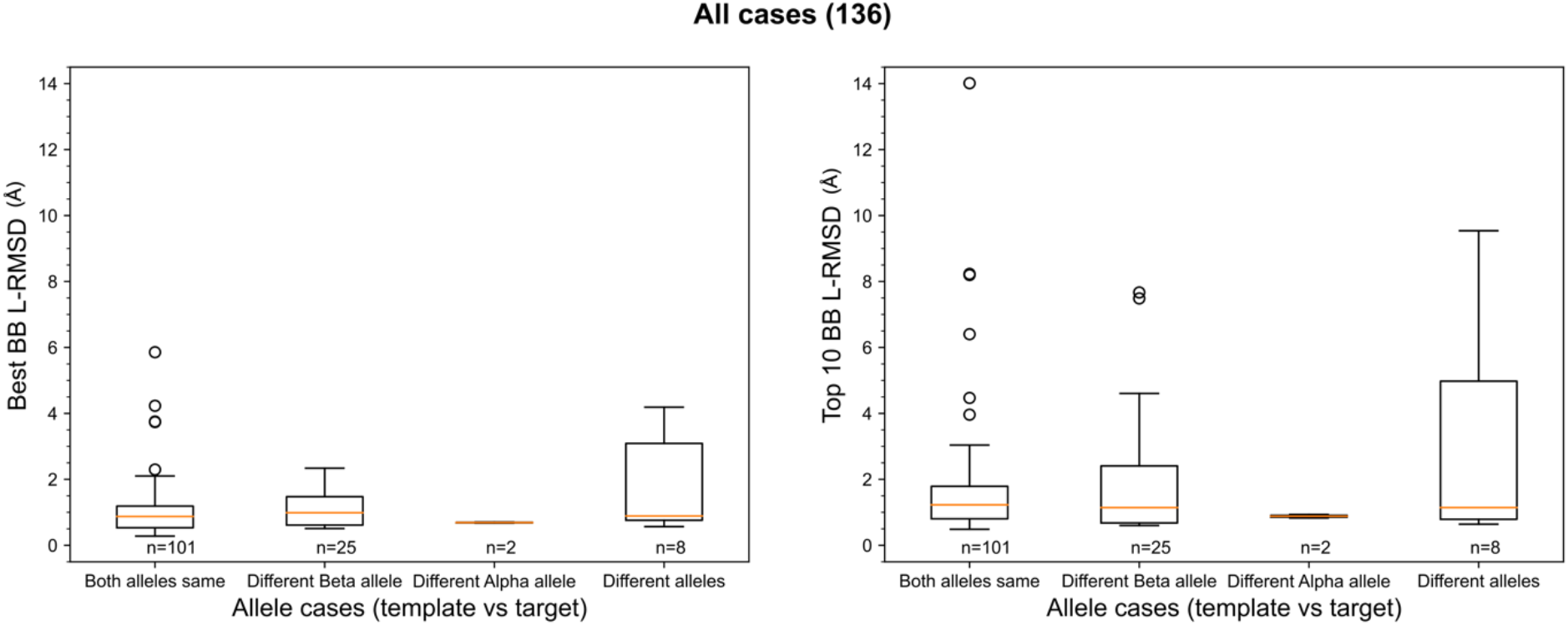
The effect of modelling with a different template allele-type and its effect on performance. Given two allele-type for each α- and β-chains of the template and target pMHC-II, 4 different scenarios are compared in each box-plot column; 1) Both allele-type is the same for template and target; 2) Only the Alpha allele-type the same;3)Only Beta allele-type the same; and 4) both chain allele-types are different in A) Best L-RMSD models and B) Top-ranked models using scoring.

### Software improvements

PANDORA v2.0 includes major improvements from the first release: Frontend (User side):

- Capability to use MHC-sequence as input instead of only allele name, leading to much broader allele coverage than version 1.0
- Addition of command-line interface for easier accessibility and bash integration
- Addition of restraints-flexible modelling mode to avoid small clashes caused by rigid restraints (see Materials and Methods)
- Improvements in the python user interface
- Easier software and database installation
- Addition of an option to remove or keep beta2-microglobulin in the generated models, as Beta2-microglobulin can be crucial or not, depending on what the models will be used for (MD, AI, manual exploration, etc.).

Backend (internal software side):

- BLAST-based template selection instead of allele-name based template selection
- Addition of a reference sequence database for allele names and MHC sequence automatic retrieval
- The allele name is now be automatically retrieved with BLAST when only the sequence is provided
- Improvement in the MHC-II template parsing to prevent multiple structures from being discarded or from missing the allele name.
- Addition of parallelization and minor optimization improvements for the template set generation, drastically increasing its speed

## Discussion

PANDORA is a 3D modelling software for both pMHC-I and -II. Here we evaluated the performance of PANDORA on reliably generating peptide conformations binding to MHC class II complexes alongside the software improvements. We applied homology-driven restraint-based modelling to reduce the computational time during sampling (3.75 min/case on one CPU core). The proposed method was tested on 136 complexes, making it the largest modeling effort of pMHC-II complexes to date. Our results show that PANDORA was able to effectively model these complexes, achieving an 85% success rate (L-RMSD < 2 Å) for the top 5 ranked models in the entire template set and generating particularly high-quality peptide core conformations.

PANDORA outperforms pDock (32) and AlphaFold (39) regarding computational time and core L-RMSD values. PANDORA incorporates domain knowledge into the modeling. In contrast, AlphaFold is a general protein structure prediction method that relies on sequence conservation information, and conservation on the peptide side has little or no bearing on this binding. Our comparison shows that not all AlphaFold-generated pMHC-II conformations have the correct anchor positions. AlphaFold’s higher computational cost is a major impediment to model millions of pMHCs, whereas PANDORA is a more practical choice.

PANDORA has the following unique features: 1) Fast: enabling high-throughput modeling of 3D pMHC-II complexes; 2) Reliable: generating low-energy models; 3) Efficient: With the use of anchor distance restraints, it to work on both MHC-I and MHC-II; 4) Template availability: providing an extensively cleaned template database of pMHC complexes, valuable for reliable homology modeling; 5) Highly Modular: It is easy to customize or extend.

PANDORA has a user-friendly interface allowing users to incorporate new configurations such as 1) more extensive sampling (especially with longer peptides); 2) specification of secondary structure restraints (23% of benchmark cases formed beta-strand PFR, Supplementary Fig. S5D); 3) fully fixed mode vs. flexible mode for the core conformation; 4) Manually defining the anchor residues; 5) Possibility of changing to other anchor predictor software. Its highly modular framework (Supplementary Fig. S6) facilitates future community-wide development.

Knowledge of the peptide binding core is required to generate the pMHC-II complex structure. When the user doesn’t input the anchor residues’ position, PANDORA currently relies on NetMHCIIpan-4.0 as an anchor predictor (36). This software has a limited, yet large, set of available MHC alleles to utilize, and it can sometimes fail to predict the correct binding core (Supplementary Figure S4). Using an anchor predictor relying on structural and physics-based data could overcome these limitations for the pMHC anchor prediction, allowing for more accurate, pan-allelic anchor predictions.

PFR can influence TCR interactions (46–49); introducing a modeling program to generate credible PFR conformations is an important step forward. It is important to note that a singular X-ray structure exclusively depicts only one snapshot of the complex conformation. This implies that a method could generate a possible PFR conformation that is not currently cataloged in the PDB but holds biological significance. To address this issue, PANDORA generates an ensemble of near-native conformations (top N-ranked conformations).

Further work is needed to model the post-translational modifications (PTM) in peptides binding to MHC, which have been shown to modulate antigen presentation and recognition (50,51) and, moreover, PTMs on peptides increase the vast number of possible pMHC combinations. PTMs have a structural impact on the stability of pMHC complexes and the consequent modulations of immune responses (52). Although it has not yet been extensively evaluated within our framework, we recognize its potential benefits for the field and remain committed to conducting additional research and possibly incorporating this method into our future research.

In conclusion, the ability of PANDORA to generate high-quality peptide conformations within the MHC-II binding groove lends great reliability to the models employed for analyzing molecular interactions at the atomic level. Due to PANDORA’s computational efficiency, initial conformations for molecular dynamics simulations can be quickly built. It is now feasible to enrich the actively accumulating wealth of pMHC binding affinity and mass spectrometry data with 3D models and aid structure-boosted artificial intelligence algorithms in identifying antigenic peptides (for example, by training the deep learning framework DeepRank on these 3D models). As such, it can be leveraged to identify cancer neoantigens or viral antigenic peptides that hold promise as vaccine candidates. It will therefore pave the way for developing novel cancer immunotherapies.

## Disclosure of Potential Conflicts of Interest

The authors declare they have no potential conflicts of interest.

### Authors’ Contributions

L. C. X., F. M. P., and D. F. M. conceived and designed the project. L. C X. coordinated and supervised the project. P.A.C.H. coordinated the project. F. M. P. and D. F. M. wrote the first draft of the manuscript. L. C X., G. R., P. A. C. H., and M. H. K. J. revised the manuscript. F. M. P., D. F. M. developed the computational approach and software. D. F. M. and G. C. contributed to the release of the Python and Conda package. D. F. M. and F. M. P. contributed to the documentation. All authors reviewed the manuscript and provided feedback.

## Acknowledgments

The authors would like to thank Giulia Crocioni, Derek van Tilborg, and Shahiel Maassen for their help with the software development.

## Data and Code Availability

The template database and package are available at DOI: 10.5281/zenodo.7318263 PANDORA is available as a Python package on conda at (27–31). The package source code and the documentation are available on GitHub at https://github.com/X-lab-3D/PANDORA.

