## supplementary_Figures for "PANDORA v2.0: Benchmarking peptide-MHC II models and software improvements"

**Supplementary Figure S1.**

Here, the effect of the similarity of the target to the template peptide on the generation of higher-quality conformations is investigated. The models with 100%peptide identity outperformed those with smaller peptide similarities. A higher-quality peptide conformation is typically observed when the template and the target are extremely similar (Supplementary Fig. S2A-B).

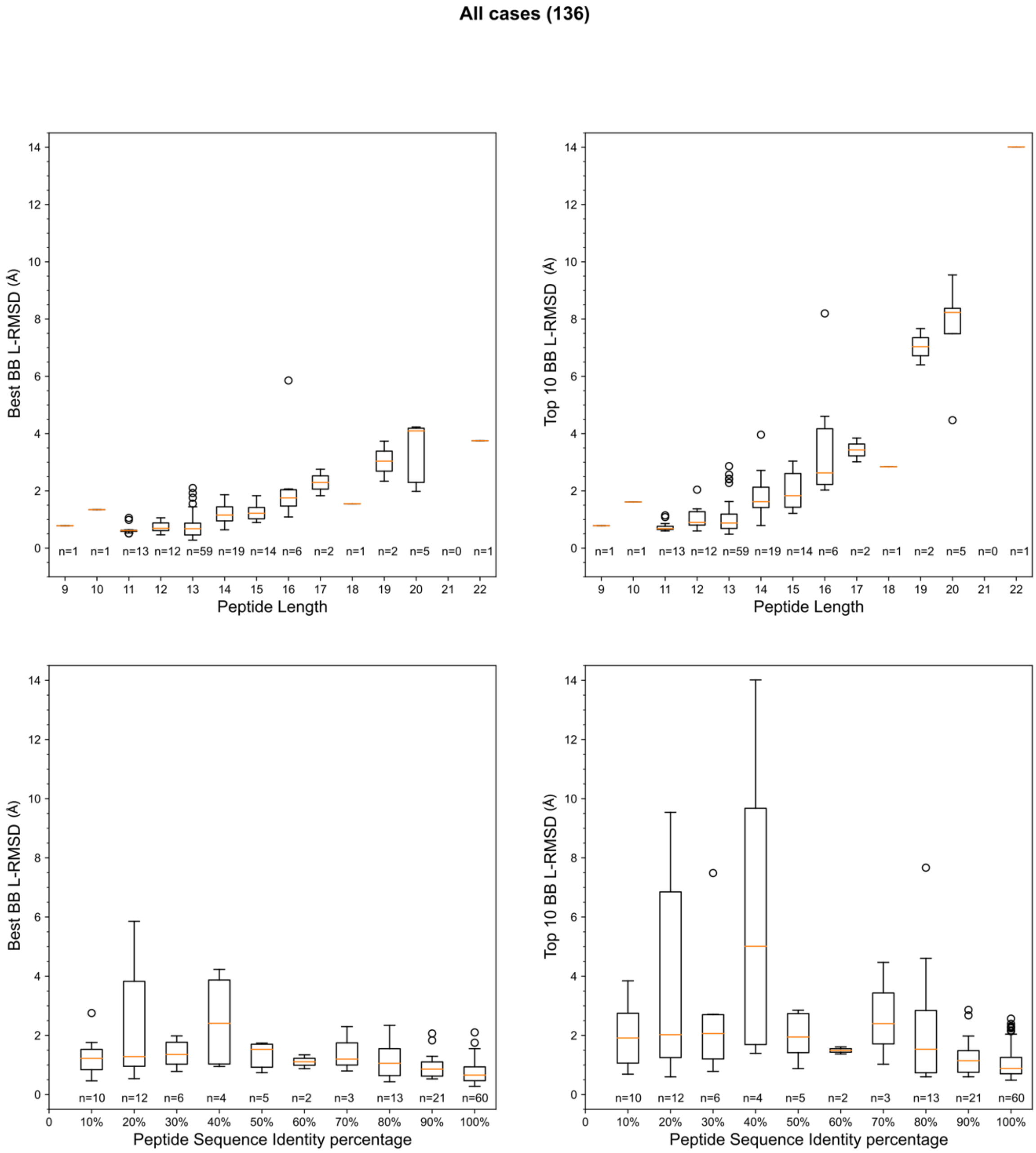

**The impacts of peptide length and peptide sequence identity on PANDORA's model performance.** The left panels depict PANDORA's sampling performance (best models generated by PANDORA among the top 20 models), while the right panels depict the scoring performance (top 10 ranked models). **(A and B)** Evaluation of PANDORA performance over different peptide lengths; **(C and D)** Evaluation of PANDORA performance over target-template peptide sequence identities.

Modeling protein-protein complexes involves two essential steps: sampling and scoring. In the first stage, known as sampling, diverse conformations are generated to explore the vast conformational space, while the subsequent stage of the scoring process involves ranking the generated models in order to identify those that are most similar to the near-native conformation. Thus, in this modelling pipeline, the generated pMHC-II models were ranked using MODELLER's internal DOPE and molpdf scores. The molpdf had the best performance by means of success-rate and hit-rate (Supplementary Fig. S1). PANDORA ranked a high-quality model (L-RMSD <1 Å) in the top 5 models, in 53% of the cases (Supplementary Fig. S2F). Approximately in 80% of the cases, the best-scored model L-RMSD is lower than 2 Å. Using a threshold of 1 Å or 2 Å as a definition of hit (model with an L-RMSD < threshold) resulted in a relatively consistent hit-rate as the number of top-N increased. This indicates that the scoring function has a marginal effect on the ratio of hits.The low L-RMSD variation between cases contributes to the consistent hit rate.

**Supplementary Figure S2.**

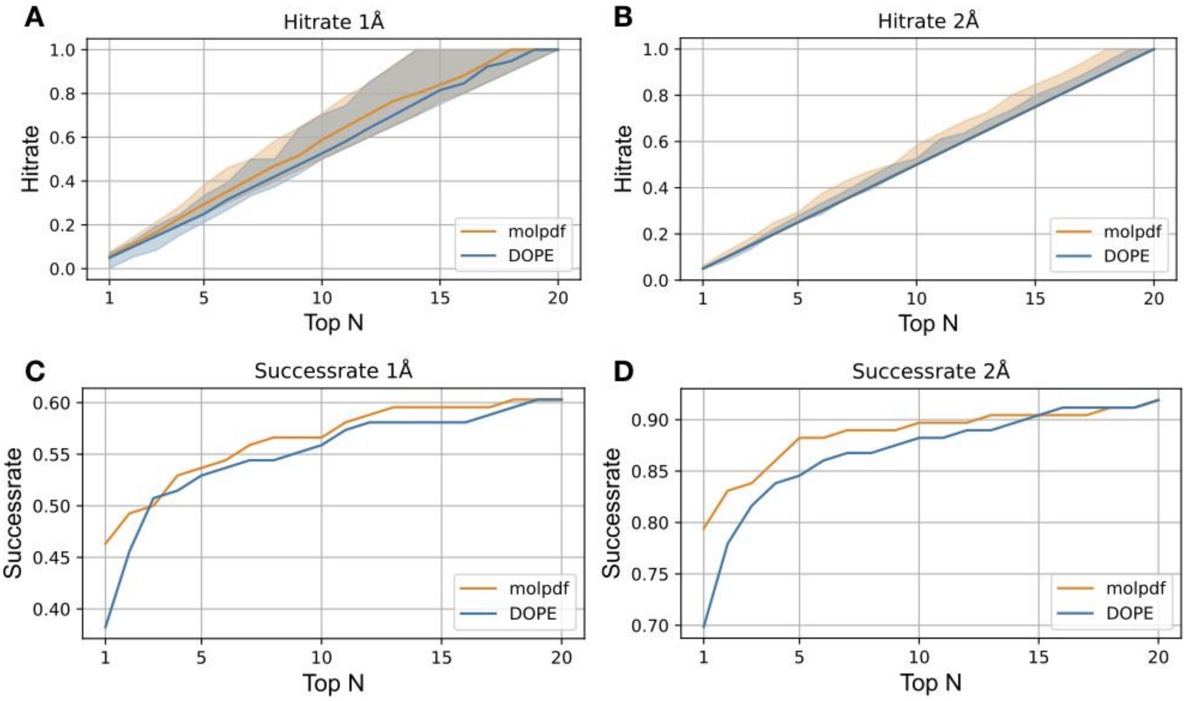

**Scoring performance using two of MODELLER's internal scoring functions, molpdf and DOPE.** A hit is defined as a model with a backbone L-RMSD ⩽ threshold. **(A, B)** Hit rate plots for two different thresholds of 1 Å and 2 Å respectively. The hit-rate shows the ratio of produced models being hits in the n best models across cases. For the hit-rate plots, the line marks the average of the hit-rates, and the shaded area marks the 25%–75% quantile interval. **(C and D)** Success rate plots for two different thresholds: 1 Å and 2 Å. Success rate measures the fraction of cases that produced at least one hit in the n best-scored models.

**Supplementary Figure S3.**

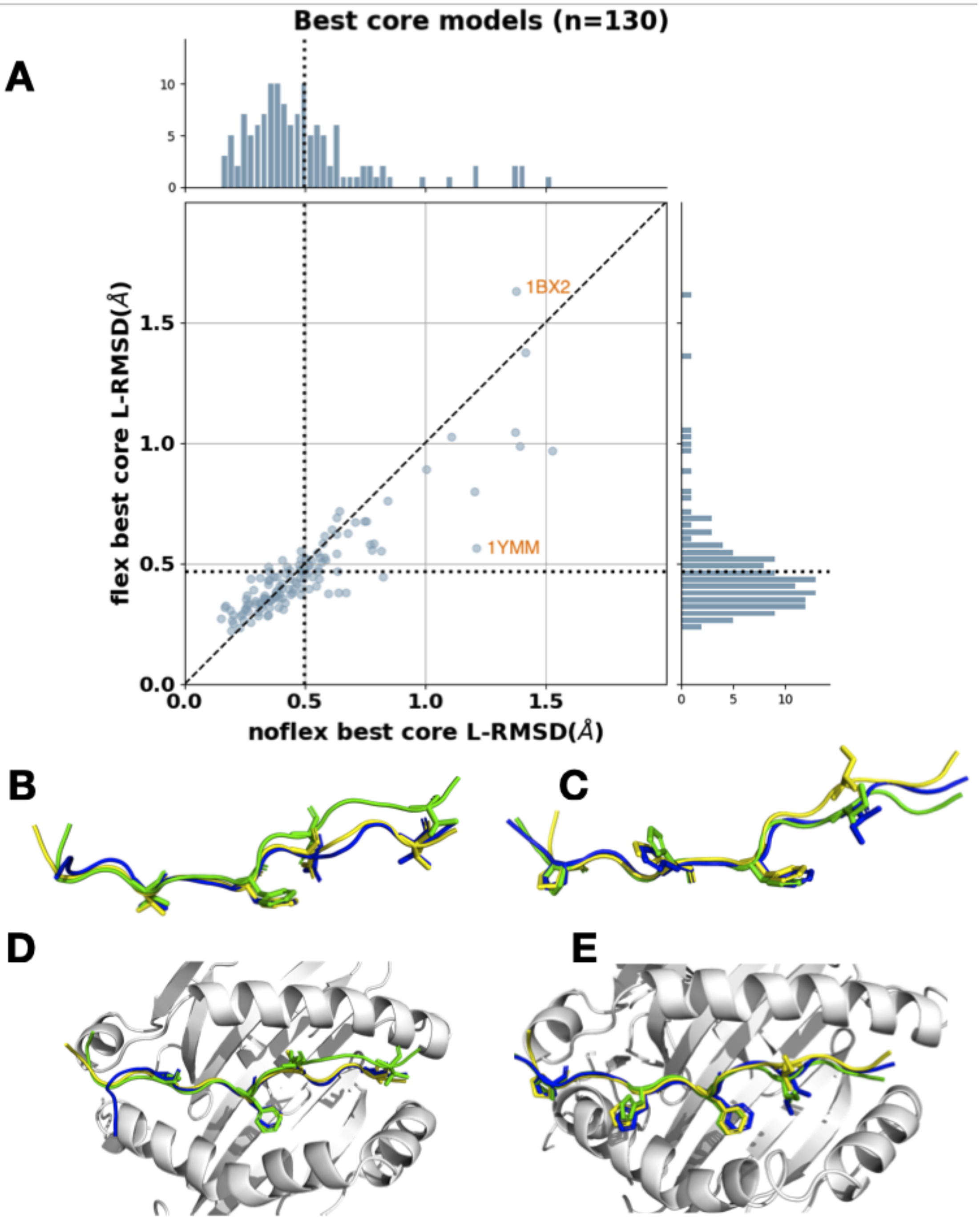


**Evaluating the two scenarios of fixed core vs. flexible core modelling. (A)** Comparison of the best core backbone L-RMSD models with both experiments. The X-axis represents the results with fixed core L-RMSD while the Y-axis represents flexible core L-RMSD. The dotted line indicates the average L-RMSD for each experiment. **(B, D)** A representative model resulting in higher core L-RMSD with the flexible experiment (fixed core L-RMSD: 1.40 Å, flexible core L-RMSD: 1.62 Å) with two different angle representations. The initial model is colored as yellow; the final PANDORA model (best-RMSD among top 5 models) is depicted as blue; and the reference structure is depicted in green (PDB ID: 1BX2) **(C, E)** A representative model resulting in lower core L-RMSD with the flexible experiment (fixed core L-RMSD: 0.56 Å, flexible core L-RMSD: 1.30 Å) with two different angle representations. The initial model is colored yellow; the final PANDORA model (best-RMSD among the top 5 models) is depicted as blue; and the reference structure is depicted in green (PDB ID: 1YMM) . Distance restraints were applied to guide the modeling process and ensure that the predicted distances between atoms were consistent with template atom distances (with a standard deviation of 0.3 Å).

**Supplementary Figure S4.**

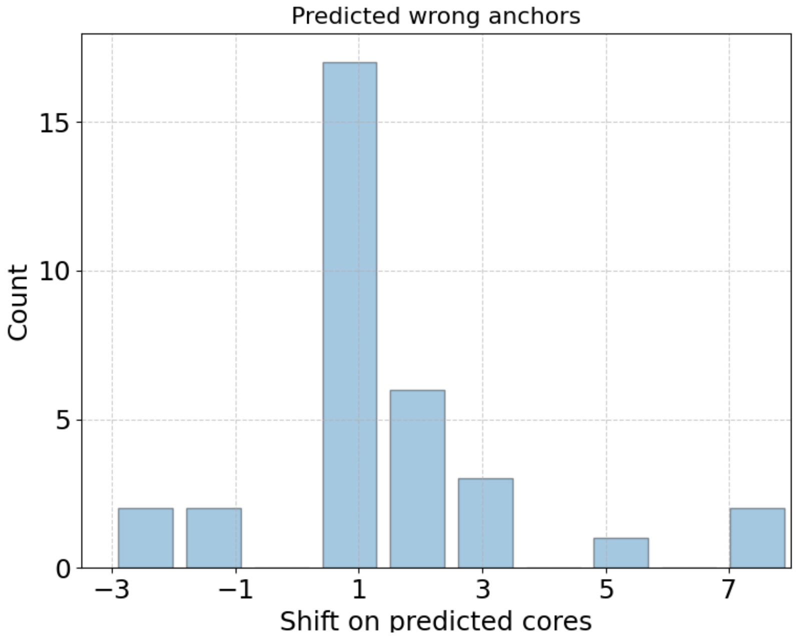


**Evaluation of the mispredicted binding cores by NetMHCIIpan-4.0 on the template set with known experimental structures**. The histogram of shifts in predicted anchors compared to real anchors.

**Conformations for PFRs and their challenges for quality assessment**

The peptide flanking regions are flexible loops that can adopt different conformations; longer PFRs exhibit more degrees of freedom. The length and structure of the PFRs affect both the binding of the peptide to MHC-II and the interaction with T-cells (Supplementary Fig. S5A). As reported (3), one indication of PFR binding to MHC-II is in the case of the formation of secondary structures while binding to MHC-II outside the binding groove (16% of the cases, Supplementary Fig. S5D). Consequently, it is essential to generate plausible structural arrangements while preserving energetically favored conformations; therefore, we demonstrate PANDORA's capacity to reproduce plausible PFR conformations.

The evaluation of the quality of generated conformations for the PFR region of the peptide is not straightforward, as some crystallized structures of pMHC-II complexes contain conformational deviations. This was caused by multiple factors, including very long PFRs (more than 8 residues at one terminus), the peptide making contact with neighboring asymmetric units (Supplementary Fig. S5C), the presence of a linker between the peptide C-terminus and MHC $\beta$-chain N-terminus (~24%, 33 out of 136), or bound to TCR, or the presence of either small molecules or crystallization cofactors close to PFRs (Supplementary Fig. S5D).

**Supplementary Figure S5.**

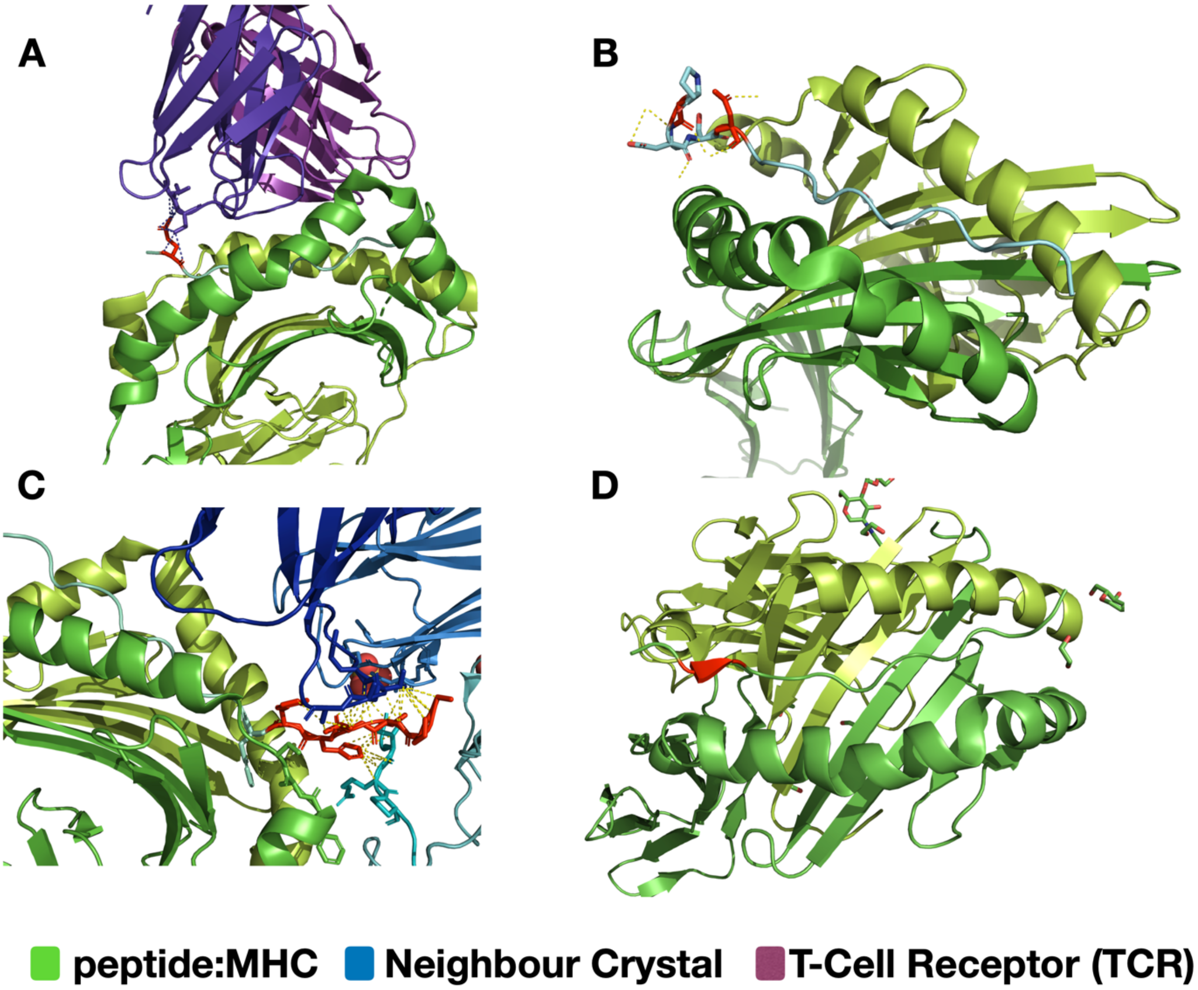


**Different folding of peptides while binding to MHC-II observed in experimental structures (A)** peptide PFR binding to TCR CDR loops **(B)** Binding of long PFR to itself **(C)** peptide PFR binding to a neighboring crystal unit **(D)** formation of a β-strand on peptide PFR along binding to MHC outside the binding groove.

**Supplementary Figure S6.**

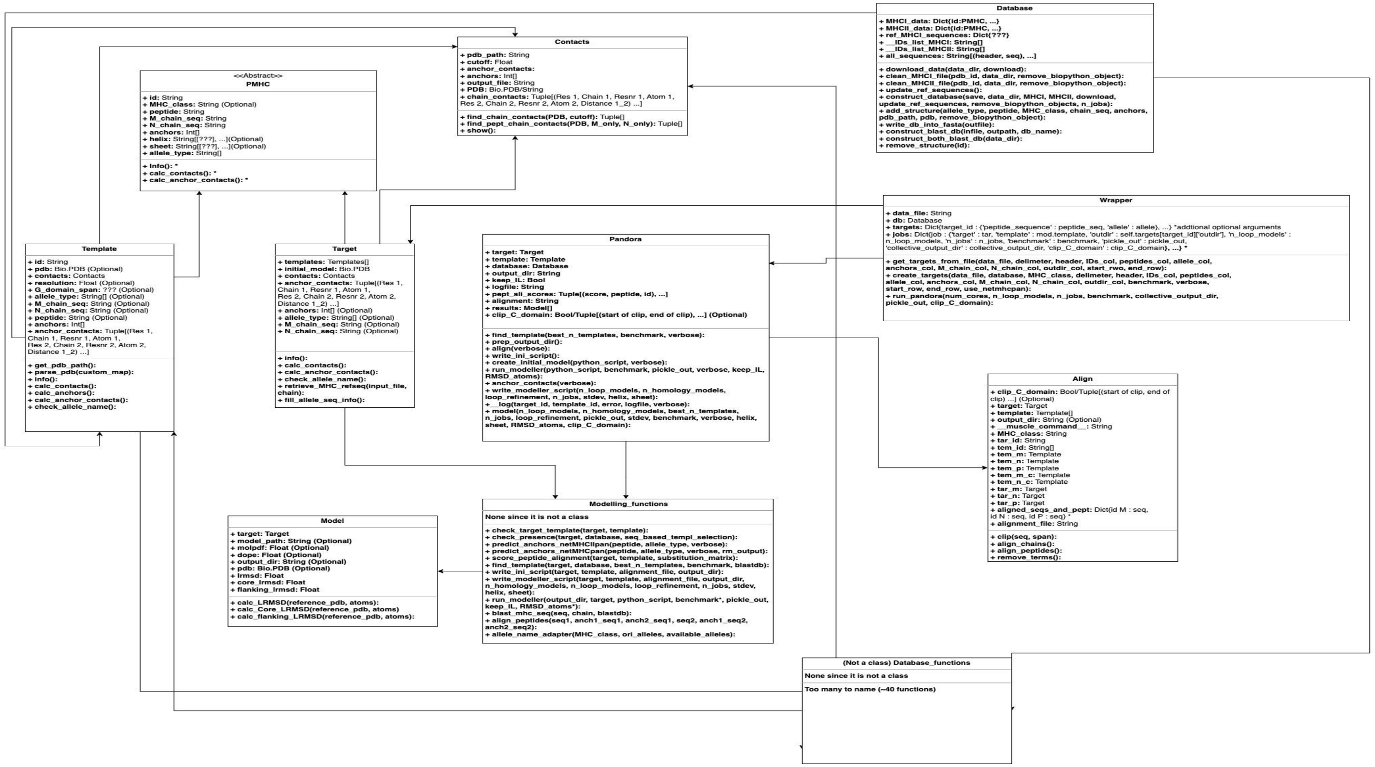


The unified modeling language (UML) Class diagram for PANDORA's modularized framework. PANDORA is highly modularized and configurable with the design for community co-development in mind.
