## supplementary_Table_1 for "PANDORA v2.0: Benchmarking peptide-MHC II models and software improvements"

**Supplementary Table 1. PANDORA cross-validation dataset.** For every case, peptide, real anchor position and IMGT alleles are reported.

| PDB ID | PEPTIDE SEQ | ANCHORS | IMGT assigned G-DOMAIN ALLELE |
|---|---|---|---|
| 1AQD | GSDWRFLRGYHQYA | 4;7;9;12 | HLA-DRA*01:02;HLA-DRA*01:01;HLA-DRB1*01:01 |
| 1BX2 | ENPVVHFFKNIVTP | 5;8;10;13 | HLA-DRA*01:02;HLA-DRA*01:01;HLA-DRB1*15:01 |
| 1D9K | GNSHRGAIEWEGIESG | 4;7;9;12 | MH2-AA*02;H2-ABk |
| 1DLH | PKYVKQNTLKLAT | 3;6;8;11 | HLA-DRA*01:01;HLA-DRA*01:02;HLA-DRB1*01:01 |
| 1ES0 | YEIAPVFVLLEYVT | 3;6;8;11 | MH2-AAd;H2-EB1b-EB24 |
| 1F3J | AMKRHGLDNYRGYS | 4;7;9;12 | MH2-AAd;H2-AB-NOD |
| 1FV1 | NPVVHFFKNIVTPRTPPPSQ | 7;10;12;15 | HLA-DRA*01:02;HLA-DRA*01:01;HLA-DRB5*01:01 |
| 1FYT | PKYVKQNTLKLAT | 3;6;8;11 | HLA-DRA*01:01;HLA-DRA*01:02;HLA-DRB1*01:01 |
| 1HQR | VHFFKNIVTP | 4;7;9 | HLA-DRA*01:02;HLA-DRA*01:01;HLA-DRB5*01:01 |
| 1HXY | PKYVKQNTLKLAT | 3;6;8;11 | HLA-DRA*01:01;HLA-DRA*01:02;HLA-DRB1*01:01 |
| 1IAK | STDYGILQINSRW | 3;6;8;11 | MH2-AA*02;H2-ABk |
| 1IAO | RGISQAVHAAHAEI | 4;7;9;12 | MH2-AAd;H2-EBu;H2-EB1b-EB24 |
| 1J8H | PKYVKQNTLKLAT | 3;6;8;11 | HLA-DRA*01:01;HLA-DRA*01:02;HLA-DRB1*04:01 |
| 1JK8 | LVEALYLVCGERGG | 3;6;8;11 | HLA-DQA1*03:02;HLA-DQA1*03:03;HLA-DQB1*03:02 |
| 1JL4 | GNSHRGAIEWEGIESG | 4;7;9;12 | MH2-AA*02;H2-ABk |
| 1JWM | PKYVKQNTLKLAT | 3;6;8;11 | HLA-DRA*01:01;HLA-DRA*01:02;HLA-DRB1*01:01 |
| 1JWS | PKYVKQNTLKLAT | 3;6;8;11 | HLA-DRA*01:01;HLA-DRA*01:02;HLA-DRB1*01:01 |
| 1JWU | PKYVKQNTLKLAT | 3;6;8;11 | HLA-DRA*01:01;HLA-DRA*01:02;HLA-DRB1*01:01 |
| 1KG0 | PKYVKQNTLKLAT | 3;6;8;11 | HLA-DRA*01:01;HLA-DRA*01:02;HLA-DRB1*01:01 |
| 1KLG | GELIGILNAAKVPAD | 4;7;9;12 | HLA-DRA*01:01;HLA-DRA*01:02;HLA-DRB1*01:01 |
| 1KLU | GELIGTLNAAKVPAD | 4;7;9;12 | HLA-DRA*01:01;HLA-DRA*01:02;HLA-DRB1*01:01 |
| 1LO5 | PKYVKQNTLKLAT | 3;6;8;11 | HLA-DRA*01:01;HLA-DRA*01:02;HLA-DRB1*01:01 |
| 1R5I | PKYVKQNTLKLAT | 3;6;8;11 | HLA-DRA*01:01;HLA-DRA*01:02;HLA-DRB1*01:01 |

| | | | |
|---|---|---|---|
| 1R5V | ADLIAYPKAATKF | 4;7;9;12 | H2-EAck;H2-EAk;MH2-AAe;H2-EB1k |
| 1R5W | ADLIAYFKAATKF | 4;7;9;12 | H2-EAk;MH2-AAe;H2-EA;H2-EAck;H2-EB1k |
| 1S9V | LQPFPQPELPY | 3;6;8;11 | HLA-DQA1*05:07;HLA-DQA1*05:05;HLA-DQA1*05:03;HLA-DQA1*05:06;HLA-DQA1*05:01;HLA-DQA1*05:08;HLA-DQB1*02:02;HLA-DQB1*02:01;HLA-DQB1*02:04 |
| 1SJE | PEVIPMFSALSEGAT | 3;6;8;11 | HLA-DRA*01:01;HLA-DRA*01:02;HLA-DRB1*01:01 |
| 1SJH | PEVIPMFSALSEG | 3;6;8;11 | HLA-DRA*01:02;HLA-DRA*01:01;HLA-DRB1*01:01 |
| 1T5W | AAYSDQATPLLLS | 3;6;8;11 | HLA-DRA*01:02;HLA-DRA*01:01;HLA-DRB1*01:01 |
| 1T5X | AAYSDQATPLLLS | 3;6;8;11 | HLA-DRA*01:01;HLA-DRA*01:02;HLA-DRB1*01:01 |
| 1U3H | SRGGASQYRPSQ | 3;6;8;11 | MH2-AAu;MH2-AA-N2W;H2-ABu |
| 1UVQ | MNLPSTKVSWAAVGGGGSLV | 3;6;8;11 | HLA-DQA1*01:02;HLA-DQB1*06:02 |
| 1YMM | ENPVVHFFKNIVTP | 5;8;10;13 | HLA-DRA*01:01;HLA-DRA*01:02;HLA-DRB1*15:01 |
| 1ZGL | VHFFKNIVTPRTPG | 4;7;9;12 | HLA-DRA*01:02;HLA-DRA*01:01;HLA-DRB5*01:01 |
| 2FSE | AGFKGEQGPKGEPG | 3;6;8;11 | HLA-DRA*01:01;HLA-DRA*01:02;H2-EBu |
| 2G9H | PKYVKQNTLKLAT | 3;6;8;11 | HLA-DRA*01:02;HLA-DRA*01:01;HLA-DRB1*01:01 |
| 2IAD | GHATQGVTAASSHE | 4;7;9;12 | MH2-AAd;H2-ABd |
| 2IAM | GELIGILNAAKVPAD | 4;7;9;12 | HLA-DRA*01:02;HLA-DRA*01:01;HLA-DRB1*01:01 |
| 2IAN | GELIGTLNAAKVPAD | 4;7;9;12 | HLA-DRA*01:02;HLA-DRA*01:01;HLA-DRB1*01:01 |
| 2ICW | PKYVKQNTLKLAT | 3;6;8;11 | HLA-DRA*01:01;HLA-DRA*01:02;HLA-DRB1*01:01 |
| 2NNA | SGEGSFQPSQENP | 3;6;8;11 | HLA-DQA1*03:03;HLA-DQA1*03:01;HLA-DQA1*03:02;HLA-DQB1*03:02 |
| 2OJE | PKYVKQNTLKLAT | 3;6;8;11 | HLA-DRA*01:01;HLA-DRA*01:02;HLA-DRB1*01:01 |
| 2PXY | RGGASQYRPSQ | 2;5;7;10 | MH2-AA-N2W;MH2-AAu;H2-ABu |
| 2Q6W | AWRSDEALPLG | 2;5;7;10 | HLA-DRA*01:01;HLA-DRA*01:02;HLA-DRB3*01:01 |
| 2SEB | AYMRADAAAGGA | 3;6;8;11 | HLA-DRA*01:02;HLA-DRA*01:01;HLA-DRB1*04:01 |

| | | | |
|---|---|---|---|
| 2Z31 | RGGASQYRPSQ | 2;5;7;10 | MH2-AAu;MH2-AA-N2W;H2-ABu |
| 3C5J | QVIILNHPGQISA | 3;6;8;11 | HLA-DRA*01:02;HLA-DRA*01:01;HLA-DRB3*03:01 |
| 3C5Z | FEAQKAKANKAVDG | 3;6;8;11 | MH2-AA;H2-AB1*01 |
| 3C60 | FEAQKAKANKAVD | 3;6;8;11 | MH2-AA;H2-AB1*01 |
| 3C6L | FEAQKAKANKAV | 3;6;8;11 | MH2-AA;H2-AB1*01 |
| 3CUP | KKMREIIGWPGGSGG | 3;6;8;11 | MH2-AAd;H2-AB-NOD |
| 3L6F | APPAYEKLSAEQSPP | 5;8;10;13 | HLA-DRA*01:02;HLA-DRA*01:01;HLA-DRB1*01:01 |
| 3LQZ | RKFHYLPFLPSTGGS | 3;6;8;11 | HLA-DPA1*01:03;HLA-DPB1*26:02 |
| 3MBE | GAMKRHGLDNYRGYSLG | 5;8;10;13 | MH2-AAd;H2-AB-NOD |
| 3PL6 | NPVVHFFKNIVTPR | 4;7;9;12 | HLA-DQA1*01:02;HLA-DQB1*05:01 |
| 3QIB | ADLIAYLKQATKG | 4;7;9;12 | H2-EAk;H2-EAck;MH2-AAe;H2-EA;H2-EB1k |
| 3QIU | ADLIAYLKQATKG | 4;7;9;12 | H2-EAk;MH2-AAe;H2-EAck;H2-EB1k |
| 3QIW | ADLIAYLEQATKG | 4;7;9;12 | MH2-AAe;H2-EAck;H2-EAk;H2-EB1k |
| 3RDT | FEAQKAKANKAVD | 3;6;8;11 | MH2-AA;H2-AB1*01 |
| 3S4S | PKYVKQNTLKLAT | 3;6;8;11 | HLA-DRA*01:01;HLA-DRB1*01:01 |
| 3S5L | PKYVKQNTLKLAT | 3;6;8;11 | HLA-DRA*01:01;HLA-DRB1*01:01 |
| 3T0E | FSWGAEGQRPGFGSGG | 3;6;8;11 | HLA-DRA*01:02;HLA-DRA*01:01;HLA-DRB1*04:01 |
| 3WEX | KVTVAFNQF | 1;4;6;9 | HLA-DPA1*02:02;HLA-DPB1*22:01;HLA-DPB1*97:01 |
| 4C56 | PKYVKQNTLKLAT | 3;6;8;11 | HLA-DRA*01:01;HLA-DRA*01:02;HLA-DRB1*01:01 |
| 4,00E+41 | GELIGILNAAKVPAD | 4;7;9;12 | HLA-DRA*01:01;HLA-DRA*01:02;HLA-DRB1*01:01 |
| 4GG6 | SGEGSFQPSQENP | 3;6;8;11 | HLA-DQA1*03:01;HLA-DQB1*03:02 |
| 4GRL | RLLMLFAKDVVSRN | 4;7;9;12 | HLA-DQA1*01:02;HLA-DQB1*05:01 |
| 4H1L | QHIRCNIPKRISA | 3;6;8;11 | HLA-DRA*01:02;HLA-DRA*01:01;HLA-DRB3*03:01 |
| 4H25 | QHIRCNIPKRIGPSKVATLVPR | 3;6;8;11 | HLA-DRA*01:01;HLA-DRA*01:02;HLA-DRB3*03:01 |
| 4H26 | QWIRVNIPKRI | 3;6;8;11 | HLA-DRA*01:02;HLA-DRA*01:01;HLA-DRB3*03:01 |

| | | | |
|---|---|---|---|
| 4I5B | VVKQNCLKLATK | 1;4;6;9 | HLA-DRA*01:01;HLA-DRA*01:02;HLA-DRB1*01:01 |
| 4IS6 | RQLYPEWTEAQRL | 3;6;8;11 | HLA-DRA*01:01;HLA-DRA*01:02;HLA-DRB1*04:01 |
| 04/May | QLVHFVRDFAQL | 3;6;8;11 | HLA-DQA1*01:02;HLA-DQB1*05:01 |
| 4OV5 | GSDARFLRGYHLYA | 4;7;9;12 | HLA-DRA*01:01;HLA-DRA*01:02;HLA-DRB1*01:01 |
| 4OZF | APQPELPYPQPGS | 2;5;7;10 | HLA-DQA1*05:06;HLA-DQA1*05:08;HLA-DQA1*05:01;HLA-DQA1*05:05;HLA-DQA1*05:07;HLA-DQA1*05:03;HLA-DQB1*02:02;HLA-DQB1*02:04;HLA-DQB1*02:01 |
| 4OZG | APQPELPYPQPGS | 2;5;7;10 | HLA-DQA1*05:06;HLA-DQA1*05:08;HLA-DQA1*05:03;HLA-DQA1*05:01;HLA-DQA1*05:07;HLA-DQA1*05:05;HLA-DQB1*02:02;HLA-DQB1*02:04;HLA-DQB1*02:01 |
| 4OZH | APQPELPYPQPG | 2;5;7;10 | HLA-DQA1*05:08;HLA-DQA1*05:01;HLA-DQA1*05:03;HLA-DQA1*05:07;HLA-DQA1*05:05;HLA-DQA1*05:06;HLA-DQB1*02:04;HLA-DQB1*02:01;HLA-DQB1*02:02 |
| 4OZI | QPFPQPELPYP | 2;5;7;10 | HLA-DQA1*05:07;HLA-DQA1*05:06;HLA-DQA1*05:03;HLA-DQA1*05:08;HLA-DQA1*05:05;HLA-DQA1*05:01;HLA-DQB1*02:02;HLA-DQB1*02:01;HLA-DQB1*02:04 |
| 4P23 | FEAQKAKANKAVD | 3;6;8;11 | MH2-AA;H2-AB1*01 |
| 4P2O | PADPLAFFSSAIKGGGGSLV | 5;8;10;13 | H2-EAk;MH2-AAe;H2-EA;H2-EAck |
| 4P2Q | ADGLAYFRSSFKGG | 4;7;9;12 | MH2-AAe;H2-EA;H2-EAck;H2-EAk;H2-EB1k |
| 4P2R | ANGVAFFLTPFKA | 4;7;9;12 | H2-EA;H2-EAck;H2-EAk;MH2-AAe;H2-EB1k |
| 4P46 | FEAQKAKANKAVD | 3;6;8;11 | MH2-AA;H2-AB1*01 |
| 4P4K | QAFWIDLFETIG | 3;6;8;11 | HLA-DPA1*01:03;HLA-DPB1*46:01;HLA-DPB1*32:01;HLA-DPB1*18:02;HLA-DPB1*41:01;HLA-DPB1*26:02;HLA-DPB1*81:01 |
| 4P4R | QAFWIDLFETIG | 3;6;8;11 | HLA-DPA1*01:03;HLA-DPB1*81:01;HLA-DPB1*32:01;HLA-DPB1*26:02;HLA-DPB1*46:01;HLA-DPB1*41:01;HLA-DPB1*18:02 |

| | | | |
|---|---|---|---|
| 4P57 | QAFWIDLFETIGGGSL | 3;6;8;11 | HLA-DPA1*01:03;HLA-DPB1*19:02;HLA-DPB1*82:01;HLA-DPB1*80:01;HLA-DPB1*94:01;HLA-DPB1*59:01;HLA-DPB1*83:01;HLA-DPB1*51:01;HLA-DPB1*77:01 |
| 4P5K | NKFDTQLFHTITGGS | 3;6;8;11 | HLA-DPA1*01:03;HLA-DPB1*46:01;HLA-DPB1*26:02;HLA-DPB1*32:01;HLA-DPB1*18:02;HLA-DPB1*41:01;HLA-DPB1*81:01 |
| 4P5M | QAYDGKDYIALKG | 3;6;8;11 | HLA-DPA1*01:03;HLA-DPB1*46:01;HLA-DPB1*41:01;HLA-DPB1*81:01;HLA-DPB1*32:01;HLA-DPB1*26:02;HLA-DPB1*18:02 |
| 4P5T | FEAQKAKANKAVD | 3;6;8;11 | MH2-AA;H2-AB1*01 |
| 4Y19 | QPLALEGSLQKRG | 3;6;8;11 | HLA-DRA*01:02;HLA-DRA*01:01;HLA-DRB1*04:26;HLA-DRB1*04:01;HLA-DRB1*04:13;HLA-DRB1*04:16;HLA-DRB1*04:64;HLA-DRB1*04:76;HLA-DRB1*04:63;HLA-DRB1*04:72;HLA-DRB1*04:38;HLA-DRB1*04:34;HLA-DRB1*04:75;HLA-DRB1*04:33 |
| 4Y1A | LQPLALEGSLQKRG | 4;7;9;12 | HLA-DRA*01:02;HLA-DRA*01:01;HLA-DRB1*04:26;HLA-DRB1*04:76;HLA-DRB1*04:72;HLA-DRB1*04:75;HLA-DRB1*04:13;HLA-DRB1*04:34;HLA-DRB1*04:63;HLA-DRB1*04:38;HLA-DRB1*04:01;HLA-DRB1*04:64;HLA-DRB1*04:16;HLA-DRB1*04:33 |
| 4Z7U | PSGEGSFQPSQENPQ | 4;7;9;12 | HLA-DQA1*03:03;HLA-DQA1*03:02;HLA-DQA1*03:01;HLA-DQB1*03:02 |
| 4Z7V | SGEGSFQPSQENP | 3;6;8;11 | HLA-DQA1*03:02;HLA-DQA1*03:01;HLA-DQA1*03:03;HLA-DQB1*03:02 |
| 4Z7W | PSGEGSFQPSQENPQ | 4;7;9;12 | HLA-DQA1*03:02;HLA-DQA1*03:01;HLA-DQA1*03:03;HLA-DQB1*03:02 |
| 5DMK | SRLGLWSRMDQL | 2;5;7;10 | MH2-AAd;H2-AB-NOD |
| 5KS9 | APSGEGSFQPSQENPQ | 5;8;10;13 | HLA-DQA1*03:01;HLA-DQA1*03:03;HLA-DQA1*03:02 |

| | | | |
|---|---|---|---|
| 5KSA | QPQQSFPEQEA | 2;5;7;10 | HLA-DQA1*05:07;HLA-DQA1*05:08;HLA-DQA1*05:06;HLA-DQA1*05:05;HLA-DQA1*05:01;HLA-DQA1*05:03 |
| 5KSB | GPQQSFPEQEA | 2;5;7;10 | HLA-DQA1*05:05;HLA-DQA1*05:08;HLA-DQA1*05:03;HLA-DQA1*05:01;HLA-DQA1*05:07;HLA-DQA1*05:06 |
| 5LAX | SKGLFRAAVPSGAS | 5;8;10;13 | HLA-DRA*01:01;HLA-DRA*01:02;HLA-DRB1*04:01 |
| 5NI9 | KRIAKAVNEKSCNC | 3;6;8;11 | HLA-DRA*01:02;HLA-DRA*01:01;HLA-DRB1*04:01 |
| 5UJT | VEELYLVAGEEGCGG | 3;6;8;11 | HLA-DQA1*03:02;HLA-DQA1*03:01;HLA-DQA1*03:03;HLA-DQB1*03:02 |
| 5V4M | GWISLWKGFSF | 3;6;8;11 | HLA-DRA*01:02;HLA-DRA*01:01;HLA-DRB1*15:01 |
| 5V4N | WISLWKGFSFGS | 1;4;6;9 | HLA-DRA*01:01;HLA-DRA*01:02;HLA-DRB1*01:01 |
| 6ATF | GVYATRSSAVRLR | 3;6;8;11 | HLA-DRA*01:01;HLA-DRA*01:02;HLA-DRB1*14:47;HLA-DRB1*14:94;HLA-DRB1*14:19;HLA-DRB1*14:02;HLA-DRB1*14:51;HLA-DRB1*14:06;HLA-DRB1*14:89;HLA-DRB1*14:13;HLA-DRB1*14:09 |
| 6BGA | ADSLSFFSSSIKRGGGSLVP | 4;7;9;12 | H2-EAk;H2-EAck;H2-EB1k |
| 6BIY | DIFERIASEASRL | 3;6;8;11 | HLA-DRA*01:01;HLA-DRA*01:02;HLA-DRB1*04:68;HLA-DRB1*04:40;HLA-DRB1*04:55;HLA-DRB1*04:42;HLA-DRB1*04:23;HLA-DRB1*04:79;HLA-DRB1*04:04;HLA-DRB1*04:56;HLA-DRB1*04:13;HLA-DRB1*04:44;HLA-DRB1*04:70 |
| 6BLQ | HLVERLYLVCGEEGAGGG | 5;8;10;13 | MH2-AAd;H2-EBu |
| 6BLX | HLVERLYLVCGGEGA | 5;8;10;13 | MH2-AAd;H2-EBu |
| 6CPL | VDRFYKTLRAEQASQE | 5;8;10;13 | HLA-DRA*01:01;HLA-DRA*01:02;HLA-DRB1*11:01 |
| 6CPN | RFYKTLRAEQASQ | 3;6;8;11 | HLA-DRA*01:01;HLA-DRA*01:02;HLA-DRB1*11:01 |
| 6CPO | RFYKTLRAEQASQ | 3;6;8;11 | HLA-DRA*01:01;HLA-DRA*01:02;HLA-DRB1*15:02 |
| 6CQJ | RFYKTLRAEQASQ | 3;6;8;11 | HLA-DRA*01:02;HLA-DRA*01:01;HLA-DRB1*01:01 |
| 6CQL | RFYKTLRAEQASQ | 3;6;8;11 | HLA-DRA*01:01;HLA-DRA*01:02;HLA-DRB1*11:01 |

| | | | |
|---|---|---|---|
| 6CQR | RFYKTLRAEQASQ | 3;6;8;11 | HLA-DRA*01:01;HLA-DRA*01:02;HLA-DRB1*01:01 |
| 6DFS | LVERLYLVCGEEGA | 4;7;9;12 | MH2-AAd |
| 6DFW | LVERLYLVCGGEG | 4;7;9;12 | MH2-AAd;HLA-DQB1*06:09;HLA-DQB1*06:05 |
| 6DFX | VEELYLVAGEEGCGGGGSL | 3;6;8;11 | HLA-DQA1*03:03;HLA-DQA1*03:01;HLA-DQA1*03:02;HLA-DQB1*03:05;HLA-DQB1*03:07 |
| 6DIG | AGNHAAGILTLGK | 3;6;8;11 | HLA-DQA1*01:02;HLA-DQB1*06:23;HLA-DQB1*06:14;HLA-DQB1*06:24;HLA-DQB1*06:33;HLA-DQB1*06:16;HLA-DQB1*06:19;HLA-DQB1*06:11;HLA-DQB1*06:10 |
| 6HBY | ARRPPLAELAALNLSGSRL | 6;9;11;14 | HLA-DRA*01:01;HLA-DRA*01:02;HLA-DRB1*01:01 |
| 6KVM | PGDSDIIRSMPEQTSEK | 6;9;11;14 | |
| 6MFF | QPFPQPEQPFP | 2;5;7;10 | HLA-DQA1*05:08;HLA-DQA1*05:05;HLA-DQA1*05:03;HLA-DQA1*05:01;HLA-DQA1*05:07;HLA-DQA1*05:06 |
| 6MFG | QPFPQPELPYP | 2;5;7;10 | HLA-DQA1*05:08;HLA-DQA1*05:07;HLA-DQA1*05:01;HLA-DQA1*05:03;HLA-DQA1*05:05;HLA-DQA1*05:06 |
| 6MKD | RVSYYGPKTSPVQ | 4;7;9;12 | MH2-AA;H2-EBu |
| 6MKR | RVSYYGPKTSPVQ | 4;7;9;12 | MH2-AA;H2-EBu |
| 6MNM | RVSYYGPKTSPVQ | 4;7;9;12 | MH2-AA;H2-EBu |
| 6MNN | RVSYYGPKTSPVQ | 4;7;9;12 | MH2-AA;H2-EBu |
| 6MNO | RVSYYGPKTSPVQ | 4;7;9;12 | MH2-AA;H2-EBu |
| 6NIX | AGFKGEQGPKGEP | 3;6;8;11 | HLA-DRA*01:02;HLA-DRA*01:01;HLA-DRB1*04:76;HLA-DRB1*04:75;HLA-DRB1*04:01;HLA-DRB1*04:13;HLA-DRB1*04:64;HLA-DRB1*04:33;HLA-DRB1*04:34;HLA-DRB1*04:63;HLA-DRB1*04:38;HLA-DRB1*04:72;HLA-DRB1*04:26;HLA-DRB1*04:16 |
| 6PX6 | APFSEQEQPVLG | 2;5;7;10 | HLA-DQA1*02:01;HLA-DQB1*02:02;HLA-DQB1*02:01 |
| 6PY2 | APFSEQEQPVLG | 2;5;7;10 | HLA-DQB1*02:02;HLA-DQB1*02:01 |

| | | | |
|---|---|---|---|
| 6U3M | QPMPMPELPYP | 2;5;7;10 | HLA-DQA1*05:07;HLA-DQA1*05:06;HLA-DQA1*05:08;HLA-DQA1*05:01;HLA-DQA1*05:03;HLA-DQA1*05:05 |
| 6U3N | APMPMPELPYP | 2;5;7;10 | HLA-DQA1*05:06;HLA-DQA1*05:05;HLA-DQA1*05:03;HLA-DQA1*05:07;HLA-DQA1*05:01;HLA-DQA1*05:08 |
| 6U3O | AVVQSELPYPEGS | 3;6;8;11 | HLA-DQA1*05:08;HLA-DQA1*05:05;HLA-DQA1*05:03;HLA-DQA1*05:01;HLA-DQA1*05:07;HLA-DQA1*05:06 |
| 6XP6 | AAPQPELPYPQPG | 3;6;8;11 | HLA-DQA1*0501;HLA-DQA1*0503;HLA-DQA1*0505;HLA-DQA1*0506;HLA-DQA1*0508;HLA-DQA1*0507;HLA-DQB1*0201;HLA-DQB1*0202 |
| 7KEI | ERNAGSGIIISDGGGGSLVP | 3;6;8;11 | HLA-DQA1*0102; HLA-DQB1*0602 |
| 7N19 | GGIGSDNKVTRRG | 3;6;8;11 | HLA-DRA*01:02;HLA-DRA*01:01;HLA-DRB1*03:01 |
