## supplementary_Table_2 for "PANDORA v2.0: Benchmarking peptide-MHC II models and software improvements"

**Supplementary Table 2. L-RMSD values of PANDORA's experments for pMHC-II complexes.** Values used for Figures 3 and 5 and Supplementary Figures 1, 2, and 3 are listed.

| PDB ID | Best core Backbone L-RMSD | Best CA L-RMSD | Best Backbone L-RMSD | Top molpdf core Backbone L-RMSD | Top molpdf CA L-RMSD | Top molpdf Backbone L-RMSD |
|---|---|---|---|---|---|---|
| 1U3H | 0.404 | 0.322 | 0.858 | 0.404 | 0.322 | 0.914 |
| 5NI9 | 0.492 | 0.675 | 0.983 | 0.495 | 0.758 | 1.07 |
| 1JWM | 0.229 | 0.391 | 0.468 | 0.232 | 0.687 | 0.83 |
| 1UVQ | 0.634 | 4.002 | 3.898 | 0.639 | 10.262 | 10.515 |
| 4E41 | 0.345 | 1.084 | 1.16 | 0.347 | 1.656 | 1.503 |
| 4P2R | 0.345 | 1.296 | 1.344 | 0.348 | 1.296 | 1.416 |
| 5KSA | 0.436 | 0.588 | 0.586 | 0.439 | 0.665 | 0.682 |
| 6DFX | 0.267 | 2.223 | 2.27 | 0.267 | 11.044 | 11.396 |
| 6U3M | 0.526 | 0.485 | 0.595 | 0.528 | 0.547 | 0.831 |
| 1YMM | 1.213 | 1.449 | 1.433 | 1.214 | 2.792 | 2.742 |
| 3C60 | 0.292 | 0.598 | 0.74 | 0.293 | 0.738 | 0.806 |
| 4P4K | 0.48 | 0.408 | 0.625 | 0.482 | 0.408 | 0.896 |
| 5LAX | 0.638 | 0.846 | 0.888 | 0.646 | 1.622 | 1.561 |
| 6DIG | 0.415 | 0.554 | 0.86 | 0.418 | 0.847 | 0.914 |
| 6U3N | 0.35 | 0.319 | 0.575 | 0.354 | 0.327 | 0.623 |
| 4P46 | 0.285 | 0.53 | 0.578 | 0.286 | 0.75 | 0.789 |
| 6HBY | 0.39 | 3.222 | 3.111 | 0.393 | 7.375 | 7.382 |
| 1JWS | 0.242 | 0.375 | 0.441 | 0.244 | 0.634 | 0.82 |
| 3C6L | 0.476 | 0.498 | 0.562 | 0.477 | 0.553 | 0.684 |
| 4GG6 | 0.425 | 0.659 | 0.737 | 0.43 | 0.768 | 0.751 |
| 5V4M | 1.109 | 0.979 | 1.023 | 1.109 | 0.983 | 1.037 |

| | | | | | | |
|---|---|---|---|---|---|---|
| 6DFW | 0.63 | 0.594 | 0.687 | 0.632 | 0.695 | 0.769 |
| 1JWU | 0.194 | 0.324 | 0.423 | 0.196 | 0.657 | 0.788 |
| 1ZGL | 1.415 | 2.049 | 1.976 | 1.422 | 2.465 | 2.206 |
| 2FSE | 1.003 | 1.091 | 1.274 | 1.005 | 1.091 | 1.274 |
| 3CUP | 0.633 | 1.083 | 1.304 | 0.634 | 3.597 | 3.478 |
| 3L6F | 0.516 | 1.05 | 1.301 | 0.52 | 1.054 | 1.301 |
| 4GRL | 0.48 | 0.957 | 1.092 | 0.498 | 1.346 | 1.325 |
| 4P4R | 0.496 | 0.45 | 0.614 | 0.496 | 0.459 | 0.949 |
| 5V4N | 0.815 | 1.245 | 1.445 | 0.819 | 1.504 | 1.595 |
| 6ATF | 0.559 | 0.516 | 0.508 | 0.563 | 0.542 | 0.673 |
| 6KVM | 0.633 | 3.003 | 2.847 | 0.65 | 4.216 | 4.148 |
| 6U3O | 0.708 | 1.093 | 1.453 | 0.715 | 2.295 | 2.335 |
| 1KG0 | 0.263 | 0.317 | 0.336 | 0.265 | 0.434 | 0.439 |
| 4H1L | 0.44 | 0.773 | 0.826 | 0.454 | 0.888 | 1.021 |
| 6XP6 | 0.431 | 0.674 | 0.617 | 0.434 | 0.674 | 0.69 |
| 7KEI | 0.668 | 4.337 | 4.322 | 0.684 | 7.286 | 7.416 |
| 2G9H | 0.291 | 0.341 | 0.419 | 0.293 | 0.464 | 0.481 |
| 1KLG | 0.306 | 1.026 | 1.035 | 0.307 | 1.241 | 1.267 |
| 3LQZ | 0.598 | 1.769 | 1.756 | 0.603 | 2.595 | 2.496 |
| 4H25 | 0.549 | 3.98 | 4.052 | 0.551 | 21.205 | 21.463 |
| 6BGA | 0.473 | 5.196 | 5.158 | 0.476 | 11.361 | 11.498 |
| 7N19 | 0.548 | 0.696 | 0.699 | 0.549 | 0.715 | 0.705 |
| 1KLU | 0.341 | 0.872 | 1.039 | 0.345 | 1.294 | 1.417 |
| 2IAD | 0.485 | 1.776 | 1.816 | 0.487 | 2.351 | 2.241 |
| 4H26 | 0.556 | 0.284 | 0.528 | 0.559 | 0.314 | 0.683 |
| 1AQD | 0.275 | 0.82 | 0.854 | 0.285 | 1.207 | 1.111 |
| 1LO5 | 0.393 | 0.473 | 0.49 | 0.396 | 0.664 | 0.727 |

| | | | | | | |
|---|---|---|---|---|---|---|
| 3MBE | 0.643 | 1.43 | 1.526 | 0.647 | 2.354 | 2.486 |
| 4I5B | 0.457 | 0.574 | 1.113 | 0.457 | 1.646 | 1.644 |
| 4P5K | 0.843 | 1.793 | 1.799 | 0.855 | 3.172 | 3.339 |
| 6MFF | 0.351 | 0.349 | 0.542 | 0.356 | 0.418 | 0.629 |
| 1BX2 | 1.378 | 1.982 | 1.792 | 1.379 | 2.529 | 2.381 |
| 2IAM | 0.355 | 0.909 | 0.848 | 0.355 | 1.464 | 1.406 |
| 4P57 | 0.483 | 2.947 | 2.978 | 0.485 | 5.245 | 5.613 |
| 6BIY | 0.493 | 0.698 | 0.868 | 0.495 | 0.971 | 0.982 |
| 1D9K | 0.57 | 1.733 | 1.632 | 0.572 | 2.093 | 1.947 |
| 2IAN | 0.371 | 0.738 | 1.042 | 0.371 | 1.393 | 1.362 |
| 3PL6 | 0.482 | 1.277 | 1.195 | 0.486 | 1.652 | 1.492 |
| 6MFG | 0.405 | 0.403 | 0.504 | 0.405 | 0.442 | 0.572 |
| 1DLH | 0.38 | 0.399 | 0.419 | 0.38 | 0.502 | 0.502 |
| 1R5I | 0.254 | 0.532 | 0.724 | 0.256 | 0.75 | 0.982 |
| 6BLQ | 0.169 | 2.208 | 2.07 | 0.169 | 2.208 | 2.214 |
| 1ES0 | 0.445 | 0.608 | 0.726 | 0.456 | 0.631 | 0.766 |
| 2ICW | 0.319 | 0.494 | 0.631 | 0.323 | 0.78 | 1.05 |
| 3QIB | 0.255 | 0.94 | 1.077 | 0.258 | 1.149 | 1.442 |
| 4IS6 | 0.825 | 2.073 | 2.212 | 0.829 | 2.207 | 2.378 |
| 6MKD | 0.393 | 0.486 | 0.557 | 0.396 | 0.509 | 0.615 |
| 1F3J | 0.572 | 0.979 | 0.931 | 0.584 | 1.599 | 1.63 |
| 1R5V | 0.691 | 1.159 | 1.063 | 0.692 | 1.282 | 1.375 |
| 2NNA | 0.46 | 0.713 | 0.834 | 0.462 | 1.127 | 1.202 |
| 3QIU | 0.252 | 0.713 | 0.806 | 0.253 | 0.907 | 1.003 |
| 45050 | 0.498 | 0.632 | 0.724 | 0.503 | 0.695 | 0.78 |
| 1FV1 | 1.393 | 2.302 | 2.165 | 1.4 | 3.673 | 3.508 |
| 4OV5 | 0.254 | 1.181 | 1.215 | 0.267 | 1.774 | 1.712 |

| | | | | | | |
|---|---|---|---|---|---|---|
| 4P5M | 0.779 | 0.745 | 0.898 | 0.786 | 0.898 | 0.993 |
| 6MKR | 0.305 | 0.32 | 0.364 | 0.308 | 0.38 | 0.424 |
| 6MNM | 0.339 | 0.355 | 0.422 | 0.342 | 0.355 | 0.422 |
| 1FYT | 0.375 | 0.496 | 0.519 | 0.377 | 0.536 | 0.569 |
| 1R5W | 0.747 | 0.796 | 0.927 | 0.748 | 1.136 | 1.34 |
| 6BLX | 0.583 | 0.747 | 0.87 | 0.585 | 1.721 | 1.581 |
| 1HQR | 1.526 | 1.226 | 1.32 | 1.527 | 1.519 | 1.714 |
| 3QIW | 0.166 | 0.684 | 0.869 | 0.17 | 0.829 | 0.945 |
| 4OZF | 0.39 | 0.848 | 1.06 | 0.401 | 2.381 | 2.755 |
| 4Y19 | 0.532 | 0.49 | 0.646 | 0.536 | 0.885 | 1.345 |
| 1HXY | 0.332 | 0.316 | 0.411 | 0.335 | 0.371 | 0.411 |
| 1S9V | 0.493 | 0.464 | 0.522 | 0.5 | 0.523 | 0.581 |
| 2OJE | 0.435 | 0.668 | 0.803 | 0.436 | 0.835 | 1.036 |
| 3RDT | 0.316 | 0.312 | 0.351 | 0.32 | 0.312 | 0.351 |
| 4P5T | 0.315 | 0.395 | 0.465 | 0.319 | 0.395 | 0.506 |
| 1IAK | 0.778 | 0.847 | 0.925 | 0.788 | 2.332 | 2.272 |
| 4Y1A | 0.581 | 1.061 | 1.142 | 0.585 | 1.108 | 1.175 |
| 6CPO | 1.206 | 1.45 | 1.653 | 1.206 | 1.931 | 2.136 |
| 6MNN | 0.197 | 0.403 | 0.431 | 0.197 | 0.403 | 0.431 |
| 1SJE | 0.199 | 1.816 | 1.669 | 0.201 | 2.08 | 2.021 |
| 2PXY | 0.378 | 0.307 | 0.662 | 0.38 | 0.339 | 0.924 |
| 3S4S | 0.231 | 0.265 | 0.286 | 0.233 | 0.392 | 0.722 |
| 4OZG | 0.36 | 0.849 | 0.897 | 0.367 | 0.849 | 0.897 |
| 5KS9 | 0.25 | 1.646 | 1.744 | 0.256 | 3.841 | 3.533 |
| 1SJH | 0.187 | 0.37 | 0.439 | 0.19 | 0.559 | 0.999 |
| 4OZH | 0.402 | 0.732 | 0.832 | 0.404 | 0.74 | 0.832 |
| 4Z7U | 0.347 | 1.162 | 1.189 | 0.351 | 1.646 | 1.618 |

| | | | | | | |
|---|---|---|---|---|---|---|
| 1IAO | 0.555 | 2.005 | 1.876 | 0.555 | 2.124 | 1.919 |
| 3S5L | 0.151 | 0.215 | 0.267 | 0.153 | 0.432 | 0.732 |
| 4OZI | 0.478 | 0.566 | 0.609 | 0.478 | 0.601 | 0.676 |
| 6MNO | 0.199 | 0.334 | 0.444 | 0.202 | 0.334 | 0.468 |
| 1J8H | 0.678 | 2.466 | 2.767 | 0.682 | 2.648 | 2.997 |
| 2Q6W | 0.785 | 0.883 | 1.0 | 0.786 | 0.905 | 1.178 |
| 5DMK | 0.404 | 0.347 | 0.432 | 0.406 | 1.187 | 1.278 |
| 6CPN | 0.35 | 0.383 | 0.496 | 0.351 | 0.678 | 0.92 |
| 1T5W | 0.259 | 0.398 | 0.433 | 0.262 | 0.613 | 0.583 |
| 3T0E | 1.372 | 4.937 | 5.113 | 1.378 | 7.421 | 7.651 |
| 4P23 | 0.345 | 0.558 | 0.715 | 0.349 | 0.723 | 0.931 |
| 4Z7V | 0.31 | 0.609 | 0.694 | 0.314 | 0.626 | 0.706 |
| 6CQL | 0.38 | 0.429 | 0.499 | 0.38 | 0.82 | 0.976 |
| 1T5X | 0.259 | 0.457 | 0.501 | 0.26 | 0.869 | 0.848 |
| 2SEB | 0.585 | 0.614 | 0.614 | 0.585 | 0.669 | 0.669 |
| 4Z7W | 0.376 | 1.236 | 1.279 | 0.377 | 1.968 | 1.912 |
| 6CPL | 0.24 | 1.15 | 1.324 | 0.242 | 2.214 | 2.028 |
| 6NIX | 0.77 | 1.048 | 1.068 | 0.774 | 1.048 | 1.123 |
| 1JK8 | 0.508 | 0.882 | 0.952 | 0.508 | 1.793 | 2.073 |
| 2Z31 | 0.417 | 0.37 | 0.586 | 0.417 | 0.403 | 0.726 |
| 6CQJ | 0.33 | 0.365 | 0.411 | 0.332 | 0.365 | 0.517 |
| 1JL4 | 0.611 | 2.055 | 1.986 | 0.617 | 3.619 | 3.536 |
| 3C5J | 0.517 | 0.648 | 0.758 | 0.518 | 1.537 | 1.813 |
| 3WEX | 0.753 | 0.63 | 0.753 | 0.757 | 0.636 | 0.757 |
| 4P2O | 0.502 | 4.389 | 4.631 | 0.512 | 10.441 | 10.801 |
| 6PX6 | 0.561 | 0.724 | 0.723 | 0.565 | 0.744 | 0.723 |
| 4P2Q | 0.352 | 1.177 | 1.243 | 0.354 | 2.556 | 2.684 |

| | | | | | | |
|---|---|---|---|---|---|---|
| 5KSB | 0.515 | 0.584 | 0.676 | 0.516 | 0.625 | 0.7 |
| 6CQR | 0.31 | 0.378 | 0.553 | 0.31 | 0.579 | 0.871 |
| 6PY2 | 0.451 | 0.541 | 0.619 | 0.453 | 0.974 | 0.992 |
| 3C5Z | 0.409 | 0.639 | 0.705 | 0.412 | 0.726 | 0.912 |
| 4C56 | 0.329 | 0.558 | 0.696 | 0.333 | 0.71 | 0.872 |
| 5UJT | 0.274 | 0.924 | 1.02 | 0.276 | 1.237 | 1.371 |
| 6DFS | 0.506 | 0.608 | 0.692 | 0.508 | 0.641 | 0.776 |
